# Dietary fibre hierarchical structure determines gut microbiota responses to apple (poly)phenols through metabolite-associated microbial guilds

**DOI:** 10.64898/2026.11.25.769217

**Authors:** Ivan M. Lopez-Rodulfo, Clay Swackhamer, Julia Bechtner, Fang Fang, Bruce Hamaker, Mario M. Martinez

**Affiliations:** InnograinLab, Food Technology Division, Department of Agricultural Engineering, University of Valladolid, Spain; Institute of Sustainable Processes, University of Valladolid, Paseo Prado de la Magdalena 3-5, 47011, Valladolid, Spain; Centre for Innovative Food (CiFOOD), Department of Food Science, Aarhus University, Agro Food Park 48, Aarhus N 8200, Denmark; Whistler Center for Carbohydrate Research, Department of Food Science, Purdue University, West Lafayette, Indiana 47907, USA; Department of Biological and Chemical Engineering - Industrial Biotechnology, Aarhus University, Gustav Wieds Vej 10 A, Aarhus C 8000, Denmark

**Keywords:** Plant cell walls, xyloglucan, pectin, cellulose, microbial fermentation, short-chain fatty acid, branched-chain fatty acid

## Abstract

Dietary fibres frequently coexist in the colon with vacuole-derived (poly)phenolic compounds following consumption of whole and minimally processed plant foods or combinations of dietary sources. However, whether (poly)phenols can direct microbial community composition and metabolic outputs during fibre fermentation, and whether such effects depend on fibre composition and physical assembly, remains unknown, possibly explaining inconsistent (poly)phenol clinical outcomes. Here, we investigated how dietary fibre hierarchical structural assembly and vacuole-derived (poly)phenols interact to shape gut microbial community composition and metabolite output. We developed tunable, (poly)phenol-free bacterial analogues of dicotyledonous primary cell walls, enabling comparison with compositionally matched but unstructured polysaccharide mixtures. Using a fully characterised apple (poly)phenol extract at physiologically relevant colonic concentrations and human faecal inocula, we show that (poly)phenols had minimal effects on overall microbial community composition and diversity but significantly altered microbial metabolic outputs. These changes were associated with subtle, context-dependent shifts in taxa assigned to putative saccharolytic and proteolytic guilds based on taxon–metabolite correlations. However, these effects occurred only within specific fermentation contexts characterized by low fermentability (e.g., cellulose and physically-assembled cell walls) or slowly fermentable substrates (xyloglucan), but not rapidly fermentable substrates (e.g., pectin or unstructured polysaccharide blends). Results revealed a hierarchical framework of microbial fermentation in which fibre composition defines metabolic potential, physical assembly governs substrate accessibility and fermentation dynamics, and (poly)phenols unlock utilization of recalcitrant dietary fibres through context-dependent trophic and antimicrobial (duplibiotic) effects on specialized microbial guilds, establishing fibre composition and physical assembly as key determinants of microbiota responses to dietary (poly)phenols.

**Graphical abstract:** 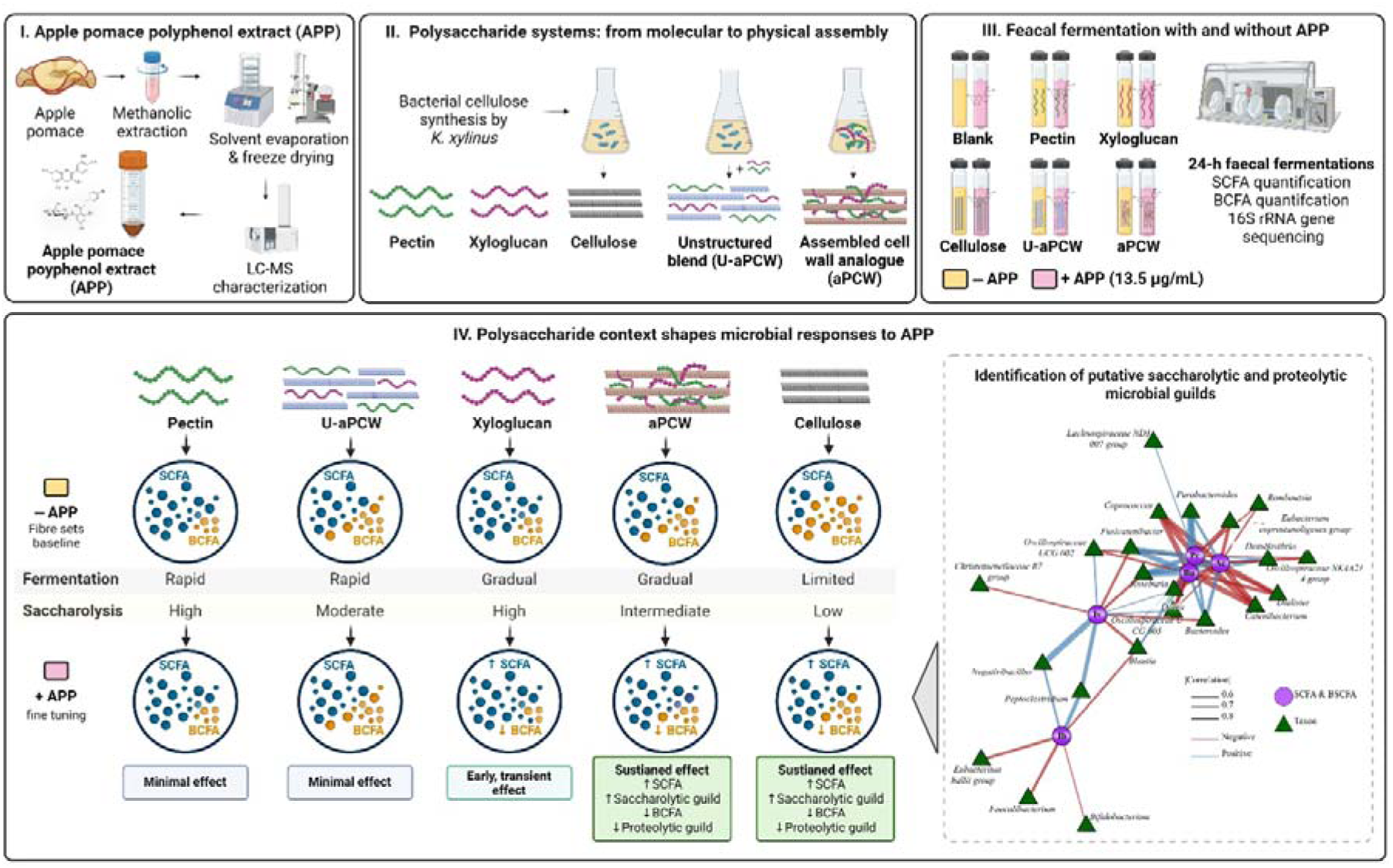

## 1. Introduction

Plant-based foods provide a diverse array of dietary fibres whose degradation by the gut microbiota is governed by structural features spanning multiple organizational scales (Deehan et al., 2020; Fu et al., 2022). At the molecular level, features such as monosaccharide composition, linkage type, branching, and degree of polymerization determine the recognition and catalytic efficiency of microbial carbohydrate-active enzymes (Hamaker & Tuncil, 2014). Beyond their molecular composition, polysaccharides assemble into higher-order structures through non-covalent interactions and polymer self-assembly. These structural arrangements determine physical properties such as porosity, hydration, surface accessibility, and spatial confinement, which ultimately influence which microbial taxa can access to fermentable substrates (Déjean et al., 2019; Diener et al., 2019; Gidley & Yakubov, 2019; Johnson et al., 2018). In whole or minimally processed foods, these polymers are embedded within three-dimensional cell wall networks rather than existing as isolated polysaccharides (Dagbasi et al., 2020; Walsh et al., 2026), limiting enzyme penetration and access to otherwise degradable glycosidic bonds (Kaur et al., 2019; von Muhlenbrock et al., 2025). In highly organized matrices, efficient deconstruction may require coordinated multienzyme systems, including cellulosome-like assemblies or other surface-associated enzymatic machineries capable of overcoming structural recalcitrance (Cantu-Jungles & Hamaker, 2020). Consequently, dietary fibre fermentability emerges from the interplay between molecular recognition and higher-order structural accessibility, highlighting the need to consider dietary fibres as hierarchical substrates whose biological functions are encoded across multiple levels of structural organization.

Notably, dietary plant cells contain vacuoles, membrane-bound intracellular compartments that store complex profiles of soluble phenolic compounds, including phenolic acids and flavonoids. Consequently, consumption of whole and minimally processed plant foods, including fruits, vegetables, and products derived from them, such as pomace, exposes the gut microbiota to intrinsically structured dietary fibre and (poly)phenol systems that are largely absent from purified dietary ingredients. In fact, the co-occurrence of dietary fibre and phenolic compounds is considered a major contributor to the health benefits associated with fruit and vegetable consumption (Bechtner et al., 2025; Cheng et al., 2025; Padayachee et al., 2017). During mastication and food processing, disruption of plant cellular structures releases vacuolar phenolic compounds while preserving varying degrees of cell wall organization. As many dietary (poly)phenols exhibit limited bioaccessibility and absorption (Lopez-Rodulfo et al., 2024; X. Zhang et al., 2020), substantial amounts reach the colon together with intrinsic dietary fibre, where they can interact simultaneously. Soluble plant-derived (poly)phenols may also be consumed independently of the fibre matrix, for example in beverages, and may interact with the gut microbiota alongside dietary fibre supplied by co-ingested foods. Beyond serving as microbial substrates, phenolic compounds are increasingly recognized as modulators of the gut microbiota, simultaneously stimulating beneficial taxa while inhibiting undesirable ones, a concept termed the duplibiotic effect (Rodríguez-Daza et al., 2021). Because many gut taxa share fermentation functions irrespective of phylogeny, such effects may be better captured at the level of microbial guilds, i.e., organisms that exploit or produce a shared key resource, with each member’s contribution depending on environmental context (Rivas-Santisteban et al., 2024).

Despite early interest in fibre–(poly)phenol interactions (Aprikian et al., 2003; Renard et al., 2001; Zduńczyk et al., 2006), clinical studies have reported inconsistent health outcomes (García-Cordero et al., 2023; C. Hou et al., 2024; Karl et al., 2025), highlighting a critical knowledge gap in how dietary fibre hierarchical structure and (poly)phenols jointly modulate gut microbiota function. Whether vacuolar (poly)phenols can be used to direct microbial community composition and metabolic outputs during dietary fibre fermentation, and whether such effects depend on the hierarchical structural assembly of the fibre, remains unknown.

In this study, we investigated how dietary fibre hierarchical structural assembly and vacuole-derived (poly)phenols interact to shape gut microbial community composition and metabolite output. We hypothesized that fibre composition and higher-order physical assembly define the ecological context in which vacuole-derived phenolic compounds exert duplibiotic effects, collectively influencing microbial substrate utilization through enzymatic compatibility and substrate accessibility. To address these questions, we developed tunable, bacterial analogues of the cell wall (aPCW) that reproduce the hierarchical architecture of dicotyledonous primary cell walls. This model enabled direct comparison between structurally organized plant cell wall–mimicking networks and compositionally matched but non-assembled polysaccharide mixtures. We further employed a fully characterized extract of vacuole-derived apple (poly)phenols at physiologically relevant colonic concentrations and investigated their effects on gut microbial community composition, diversity, and SCFA and branched-chain SCFAs (BCFAs) metabolites during dietary fibre *in vitro* fermentation with pooled human faecal inocula. SCFA and BCFA, as end-products of carbohydrate and protein fermentation, respectively, were used as the key resources defining putative saccharolytic and proteolytic guilds. Our findings reveal a hierarchical framework of microbial fermentation in which fibre composition defines metabolic potential, physical assembly governs substrate accessibility and fermentation dynamics, and vacuole-derived phenolic compounds exert context-dependent duplibiotic effects that coincide with shifts in putative saccharolytic and proteolytic microbial guilds.

## 2. Materials and methods

### 2.1. Generation and characterization of bacterial cellulose and structured primary cell wall analogues from dicotyledonous plants

Bacterial cellulose hydrogels (cellulose) were produced using *Komagataeibacter xylinus* (ATCC-53524) following a previously reported method (Mikkelsen et al., 2020). To better reproduce the composition and hierarchical organization of primary dicotyledonous cell walls, three-component bacterial hydrogels (aPCW) were generated by *K. xylinus* in the presence of pectin and xyloglucan, following Mikkelsen et al. (2020), with modifications to this protocol and characterization as described by Bechtner et al. (2025). Prior to characterization and in vitro fermentation experiments, both cellulose and aPCW hydrogels were freeze-dried and ball-milled into fine powders.

aPCW exhibited 14.2 ± 2.0 % (w/w) pectin (quantified by the m-hydroxydiphenyl assay and expressed in galacturonic acid equivalents) and 9.8 ± 0.7 % (w/w) xyloglucan, determined after double hydrolysis in 2 M methanolic HCl for 5 h and 100 °C followed by 0.5 M TFA for 1 h and 121 °C and HPAEC–PAD analysis on a CarboPac PA-1 column (ICS-6000, Thermo Fisher). The hierarchical supramolecular architecture of aPCW was confirmed by CP-MAS NMR, while the homogeneous embedding of cellulose fibrils within a continuous pectin–xyloglucan matrix was further verified by scanning electron microscopy (Bechtner et al., 2025; Fang et al., 2026).

### 2.2. Preparation of an extract of vacuole-derived phenolic compounds from apple pomace

A free-methanolic extract of vacuolar apple pomace polyphenols (APP) was prepared as previously described (Lopez-Rodulfo et al., 2024, 2026). Briefly, apple pomace obtained by cold pressing (5,000 psi; stem and carpel excluded) was freeze-dried, milled, and extracted four times with methanol–formic acid–water (80:1:19, v/v/v) under agitation (2 h, 4 °C, darkness), followed by a final 12 h extraction. Supernatants were pooled, concentrated by rotary evaporation (20 mbar, 35 °C), freeze-dried, and stored at −70 °C. The polyphenol profile, previously characterized by UHPLC-ESI-QTOF-MS, comprised flavonols (22%), polymeric flavanols, DP 5–7 (5%), flavanols, DP 1–4 (15%), dihydrochalcones (10%), hydroxycinnamic acids (22%), and hydroxybenzoic acids (27%) (Lopez-Rodulfo et al., 2024).

### 2.3. Substrate design and experimental treatments

The following substrates were prepared: (1) cellulose, (2) aPCW; (3) an unstructured blend of cellulose, xyloglucan, and pectin at the same mass ratio (76:14:10) as that constituted within aPCW (U-aPCW); (4) pectin and (5) xyloglucan. Additional controls included Blank (no substrate), APP alone, and fructooligosaccharide (FOS) as a positive control.

### 2.4. In vitro faecal batch fermentations, microbiota and fermentation metabolite analysis

*In vitro* faecal fermentations were conducted as described previously (Rose et al., 2010). All handling was performed anaerobically (85% N□, 10% H□, 5% CO□; BactronEZ chamber, Shel Lab, USA). Substrates (50 mg dry weight) were hydrated in 4 mL reduced carbonate-phosphate buffer (0.1 M, pH 6.8), in the absence or presence of APP, added at the bioaccessible (poly)phenol concentration of digested apple pomace (13.5 µg/mL buffer; 54 µg per 50 mg fibre; 12% bioaccessibility (Lopez-Rodulfo et al., 2024, 2025)). All preparations were equilibrated overnight in the anaerobic chamber before inoculation.

Fresh faecal samples were collected from three healthy adults (two males, one female; 29–40 years, mean age 32) on habitual diets, antibiotic-free for ≥3–6 months. Written informed consent was obtained from all participants prior to sample collection, and the protocol was approved by the Institutional Review Board at Purdue University (IRB #2023-75). Samples were homogenized, diluted (1:3, w/v) in anaerobic carbonate–phosphate buffer, filtered through four layers of sterile cheesecloth, and pooled in equal proportions to generate the inoculum. Fermentations were initiated by adding 1 mL of pooled inoculum to each substrate (n = 5) and incubated at 37 °C for 24 h. Samples were collected at 0, 6, 12, and 24 h for SCFA and BCFA analysis, with additional aliquots at 0 and 24 h reserved for DNA extraction. All samples were stored at −80 °C until analysis.

### 2.5. Fermentation metabolite analysis

Analysis of SCFA and BCFA was performed through GC-FID (Tuncil et al., 2017) including acetate, propionate, butyrate, isobutyrate, and isovalerate as external standards, and 4-Methylvaleric acid as internal standard. Supernatants from fermentation were collected for quantification with a gas chromatograph (GC-FID 7890A, Agilent Technologies, USA) equipped with a fused silica capillary column (Nukon™; Supelco No. 40369-03A, USA). Chromatographic conditions were: injection volume 4 μL, injector temperature 230 °C; detector temperature 230 °C; initial oven temperature 100 °C; temperature ramp of 8 °C/min to 200 °C with a 3-min hold; and helium as carrier gas (0.75 mL/min).

### 2.6. DNA isolation, microbiota profiling, by 16S rRNA gene sequencing and data analysis

Frozen fermentation samples collected at 0 and 24 h were thawed and centrifuged at 13,000 rpm for 10 min. Pellets were homogenized in phosphate buffer and lysed using Lysing Matrix E tubes from the FastDNA SPIN® Kit for Feces (MP Biomedicals, Santa Ana, CA, USA). Automated DNA extraction was performed using a QIAcube Connect instrument with the QIAamp PowerFecal Pro DNA QIAcube Kit (Qiagen, Germantown, MD, USA) according to the manufacturer’s instructions. DNA concentration was determined using a NanoDrop 2000c spectrophotometer (Thermo Fisher Scientific, Pittsburgh, PA, USA).

For 16S rRNA gene analysis, the V4 region was amplified using primers 515F (5′-GTGCCAGCMGCCGCGGTAA-3′) and 806R (5′-GGACTACHVGGGTWTCTAAT-3′). PCR amplification was performed using Phusion High-Fidelity PCR Master Mix (Thermo Fisher Scientific, USA). Amplicon concentration and purity were assessed using a NanoPhotometer NP60. Sequencing libraries were prepared using the Illumina DNA Prep kit, followed by paired-end sequencing (2 × 250 bp) on an Illumina MiSeq platform. DNA amplification, library preparation, and sequencing were conducted at the Purdue Genomics Core Facility (West Lafayette, IN, USA).

Processing and analysing the 16S rRNA gene sequence data were conducted using methods established in previous studies (Johnson et al., 2019). Briefly, raw sequence data were imported into the CLC Genomics Workbench (QIAGEN, Redwood City, CA, USA) for demultiplexing, quality trimming (Q20), and removal of reads shorter than 200 bases. More than 30,000 high-quality reads were generated per sample Analysis of the demultiplexed, paired-end sequence files was carried out using QIIME2 2022.8. Paired-end reads were filtered, denoised, merged, and chimeras were removed using *dada2 denoise-paired* (Callahan et al., 2016), yielding exact sequence variants (ESVs). Alignment of sequences and construction of a phylogenetic tree was carried out using *phylogeny align-to-tree-mafft-fasttree*. Taxonomy was assigned to ESVs using a naïve Bayes classifier trained on the Greengenes 13_8 99% Operational Taxonomic Units (OTUs) database (McDonald et al., 2012), which was implemented using the plugin *feature-classifier classify-sklearn* (Bokulich et al., 2018). For analysis of alpha and beta diversity, samples were first rarefied to 6,000 reads per sample using the *diversity alpha-rarefaction* plugin to reduce the effect of sequencing depth. This rarefaction depth did not influence the richness or evenness of the samples, as indicated by a plateau in the rarefaction curve of the Shannon index.

Alpha diversity metrics (Observed richness, Shannon index, Pielou’s evenness, and Chao1) were calculated, and beta diversity was assessed using Bray–Curtis dissimilarity and visualized by principal coordinates analysis (PCoA). Follow up analyses were carried out using R (v4.3.1). Low-prevalence (<10% of samples) and low-abundance taxa (mean relative abundance <0.5%) were removed prior to analysis. Data were organized using the phyloseq package for ecological analyses.

### 2.7. Statistical analysis

All fermentations were performed in parallel (n = 5) from the same pooled inoculum batch, and data are presented as mean ± SD unless otherwise stated. Statistical analyses and graphics were performed in R (v4.3.1), Python (v3.x) and OriginPro® 2023 (OriginLab, USA). Differences among substrates were tested by one-way ANOVA with Tukey’s post hoc test, and the effect of APP within each substrate and time point by Welch’s *t*-test. Differences in the APP effect (ΔAPP) between conditions were assessed by pairwise two-way ANOVA (condition × APP interaction) with Holm correction. For correlation analyses, taxonomic abundances were CLR-transformed after CZM zero replacement, and differential abundance was assessed with DESeq2 (Love et al., 2014). *P*-values from correlation and differential abundance analyses were adjusted by the Benjamini–Hochberg method. Statistical significance was set at *p* < 0.05.

## 3. Results

### 3.1. Molecular composition of dietary fibres outweighs physical assembly and the co-presence of phenolic compounds in shaping microbial community structure, diversity and taxonomic clustering

16S rRNA gene sequencing was combined with targeted SCFA and BCFA analyses across a defined panel of dietary fibre substrates. The substrate panel comprised the individual polysaccharides cellulose, pectin, and xyloglucan, together with two ternary mixtures: an analogue of dicotyledonous primary cell walls (aPCW), reproducing both composition and physical assembly, and its unstructured counterpart (U-aPCW), consisting of the same polysaccharides without higher-order organization. All substrates were fermented for 24 h with pooled human faecal inoculum in the absence or presence of APP.

Dietary fibre substrate was the primary determinant of microbial diversity and community structure after 24 h, whereas APP exerted only minor effects (**Figure 1**). Alpha diversity differed significantly among treatments (Shannon index, **Figure 1A**). Among controls, the initial inoculum, Blank, and APP-alone showed the highest diversity, while FOS showed the lowest. Among the plant cell wall polysaccharides, cellulose maintained the highest alpha diversity, pectin the lowest, and xyloglucan together with the two ternary substrates (aPCW and U-aPCW) displayed intermediate values. This inverse relationship between fibre fermentability and α-diversity likely reflects strong ecological selection, whereby fast-growing specialists rapidly capture available substrates under closed batch conditions lacking continuous substrate replenishment (Cantu-Jungles & Hamaker, 2023). Consistent with these patterns, Bray–Curtis ordination separated communities primarily according to substrate identity (**Figure 1B**). The two ternary substrates clustered closely together but remained distinct from their individual polysaccharide components, while paired ± APP treatments largely overlapped within each substrate.

**Figure 1.**
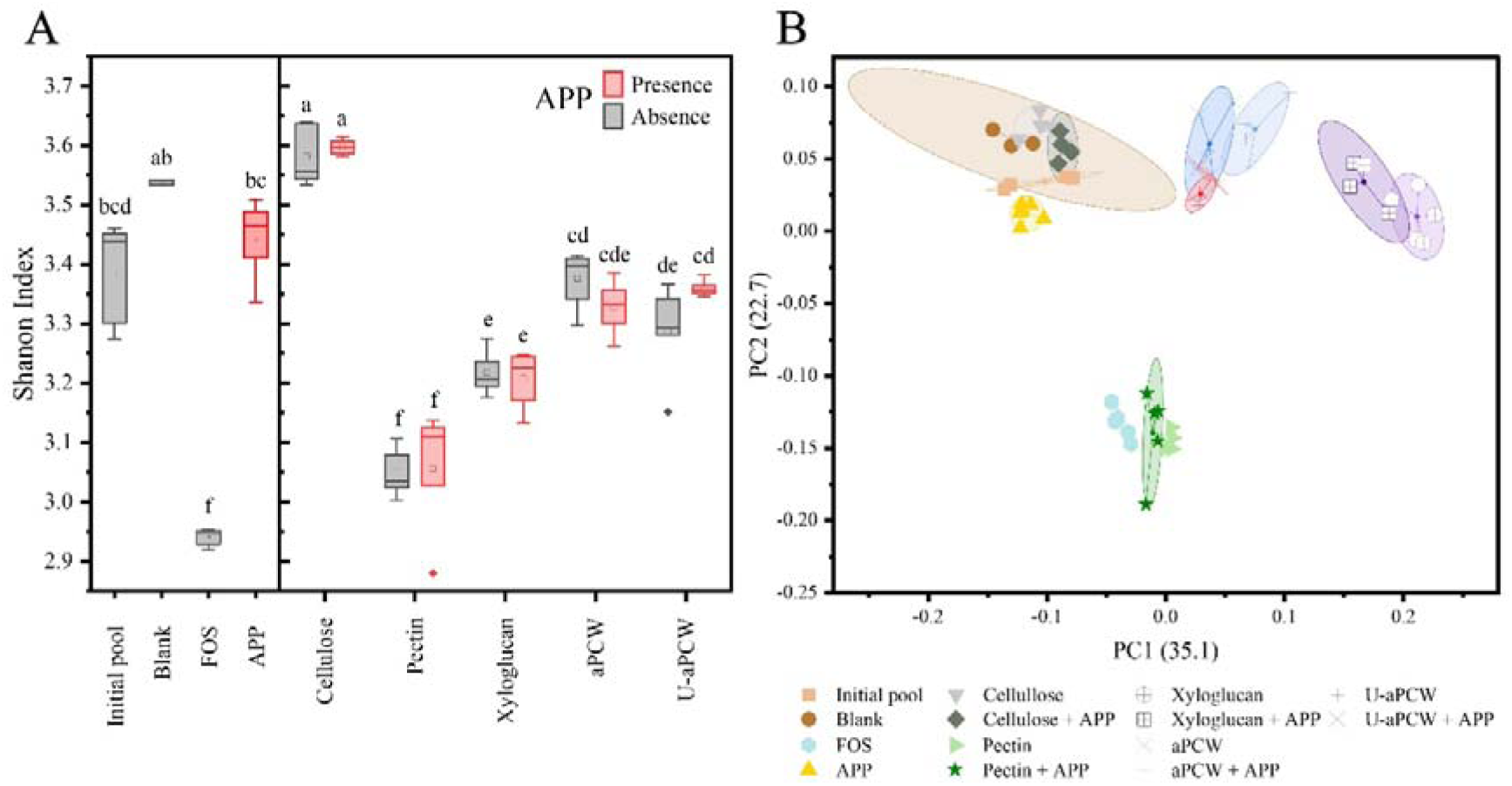
Microbial Community Structure. (A) Shannon alpha diversity of controls and substrate fermentations in the presence (red) or absence (grey) of vacuole-derived phenolic compounds (APP). Different letters indicate significant differences among substrates determined by one-way ANOVA using Tukey’s post hoc test (*p* < 0.05). (B) Principal coordinates analysis (PCoA) of microbial community structure based on Bray–Curtis dissimilarities calculated from ESV abundance profiles. Points represent individual replicates (*n* = 5).

At the phylum level (**Figure 2A**), microbial communities diverged from the initial inoculum after 24 h fermentation. Bacillota remained the dominant phylum despite decreasing from ∼84% to <79% across treatments. Highly fermentable substrates (FOS, pectin, and xyloglucan) promoted a relative expansion of Bacteroidota, whereas cellulose and the ternary substrates maintained more balanced phylum distributions. Accordingly, the Bacillota/Bacteroidota ratio (formerly Firmicutes/Bacteroidetes) decreased in highly fermentable substrates but remained higher in APP, cellulose, and the ternary substrates (**Figure S1**).

**Figure 2.**
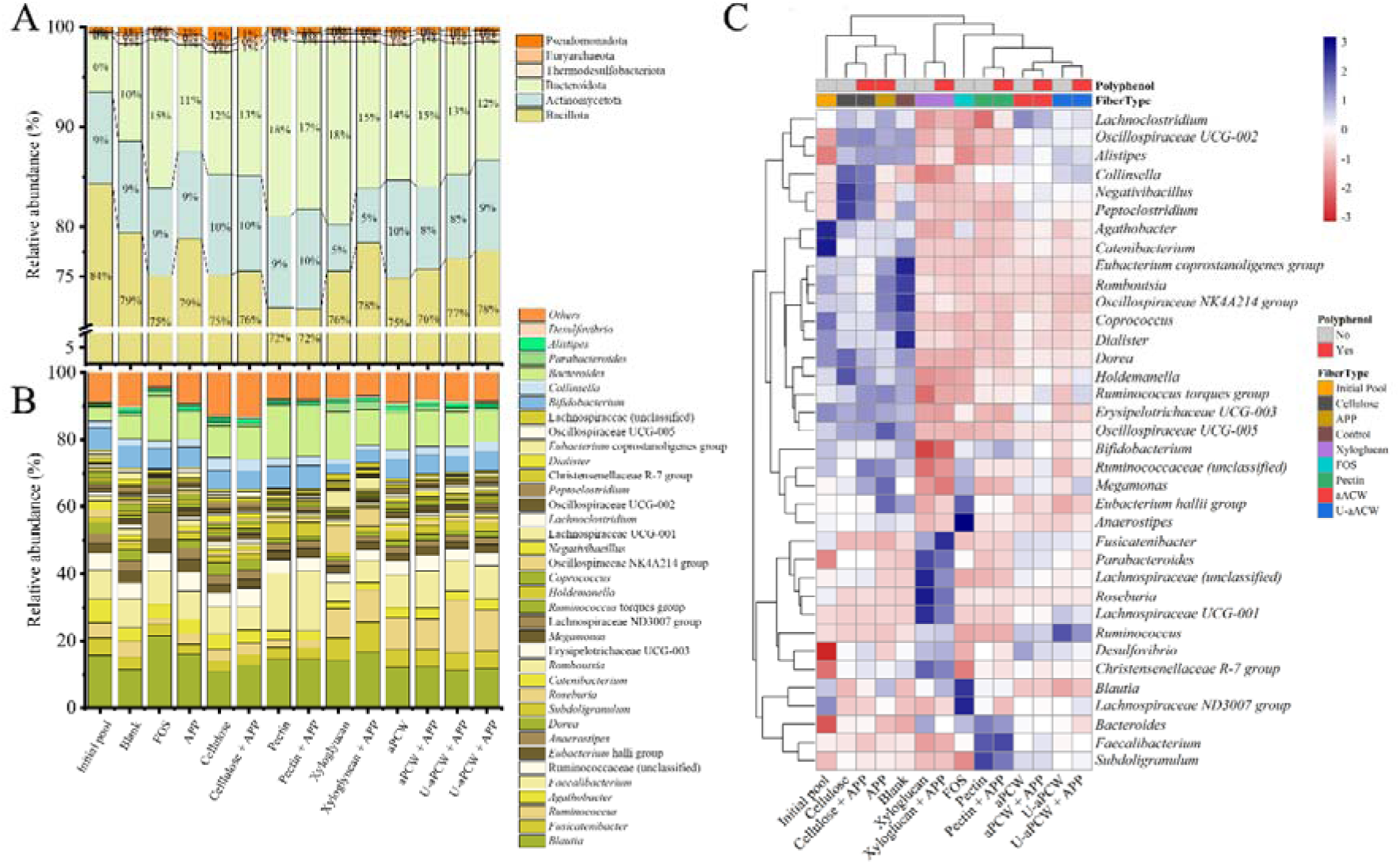
Relative abundance of dominant taxa at the phylum (A) and genus levels (B). Taxa with mean relative abundance >0.5% and prevalence ≥10% across all treatments were included; remaining taxa were grouped as others and excluded from statistical analyses. Hierarchical clustering heatmap of filtered genera across all treatments (C), including the initial pool. Clustering was performed using Euclidean distance and complete linkage. Data represent mean relative abundance across replicates (n = 5).

Genus-level profiles are shown in **Figure 2B**. Hierarchical clustering (**Figure 2C**) separated low-fermentability conditions (initial pool, blank, APP-alone, and cellulose) from fermentable fibre treatments (FOS, pectin, xyloglucan, aPCW, and U-aPCW), irrespective of APP supplementation. These data alongside taxa log2 fold-change analysis relative to the blank (**Figure S**2) further identified distinct taxonomic fingerprints for each substrate.

Cellulose grouped with low-fermentability conditions and showed an even community composition consistent with its higher Shannon index. However, relatively higher abundances of *Collinsella*, *Negativibacillus*, and *Peptoclostridium* were observed.

Within the highly fermentable-fibre group, each substrate exhibited a distinct ecological signature. FOS was characterized by enrichment of taxa such as *Anaerostipes, Blautia* and other *Lachnospiraceae* members, with a well-known saccharolytic capacity (Reichardt et al., 2018; Schwiertz et al., 2002; Zhao et al., 2024). Pectin, in contrast, enriched *Bacteroides* and *Faecalibacterium* in line with previous studies of pectinolytic communities characterized by acetogenic metabolism either through direct pectin degradation or cross-feeding interactions, accompanied by a suppression of *Peptoclostridium* (Elshahed et al., 2021; Kaoutari et al., 2013; Lopez-Siles et al., 2012; Moro Cantu-Jungles et al., 2019). Interestingly, *Subdoligranulum* was also enriched under pectin supplementation in agreement with Bulut et al. (2023). Although its potential involvement in pectin utilization has received less attention, recent genomic and experimental evidence suggests that certain strains, such as *Subdoligranulum variabile* and some *Faecalibacterium* strains may possess CAZyme-encoding genes associated with pectin utilization such as glycoside hydrolases (GHs) and carbohydrate esterases (CEs) (Solvang et al., 2025). Nevertheless, the 16S rRNA gene sequencing approach used in the present study does not allow the identification of CAZyme genes or the assignment of specific pectinolytic functions to the enriched taxa. Thus, their potential involvement in pectin fermentation remains speculative. For both FOS and pectin, the enriched taxa accounted for more than 35% of the total community.

Xyloglucan displayed a contrasting profile, enriching *Roseburia*, *Parabacteroides*, and *Bacteroides*, aligned with their known xyloglucan-degrading capacity, while the emergence of *Roseburia* may relate to its GH5 repertoire (Déjean et al., 2019; Kaoutari et al., 2013; W. Zhang et al., 2025). *Fusicatenibacter*, and other *Lachnospiraceae* members (UCG-001 and unclassified), while not recognized as xyloglucan degraders, have been reported to increase through cross-feeding on released xylose (Pihelgas et al., 2024; Zhao et al., 2024). The two ternary substrates exhibited broadly similar genus-level profiles, with moderate increases in *Lachnospiraceae* (UCG-001 and unclassified), *Roseburia*, *Ruminococcus*, and *Parabacteroides* relative to the blank. Only subtle differences were observed between aPCW and U-aPCW, including slightly higher *Lachnoclostridium* in aPCW and *Ruminococcus* in U-aPCW, without altering their shared clustering pattern.

### 3.2. Fermentation kinetics are governed by both the molecular composition and the physical assembly of dietary fibres

Microbial metabolites including SCFAs (acetate, propionate, butyrate) and BCFAs (isobutyrate and isovalerate) were quantified at 6, 12, and 24 h to elucidate how dietary fibre structural hierarchy, spanning composition to physical assembly, influences fermentability and fermentation dynamics in the absence of APP (**Figure 3**). Although SCFAs dominated total metabolite production, SCFAs and BCFAs displayed opposite responses across treatments and were therefore analysed separately **(Figure 3A–C)**.

**Figure 3.**
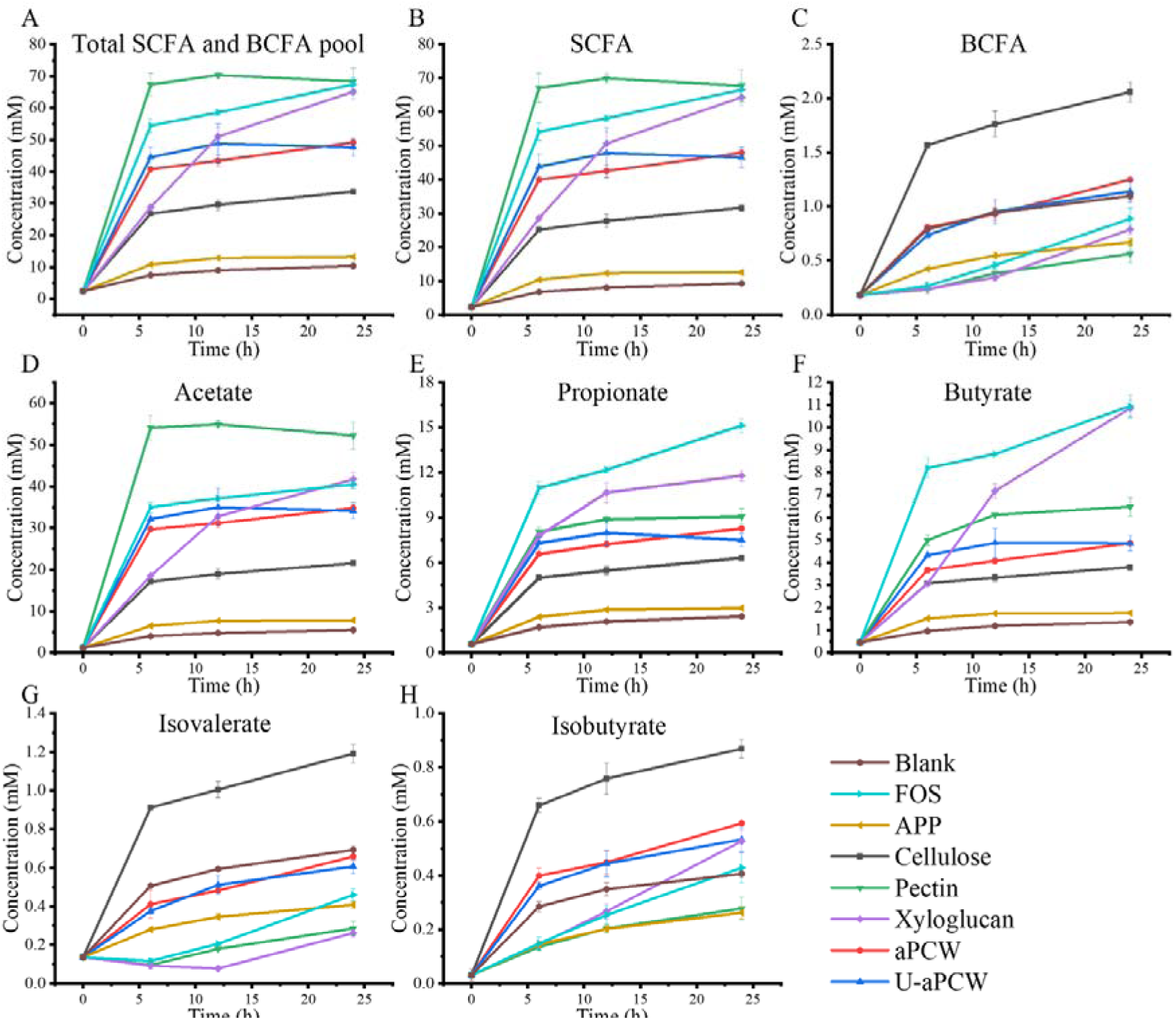
Time-resolved concentrations of total and individual SCFA and BCFA during substrate fermentations in the absence of vacuole-derived phenolic compounds (APP). Total metabolites (A). Total SCFA (B), calculated as the sum of acetate (D), propionate (E), and butyrate (F). Total BCFA (C), calculated as the sum of isovalerate (G) and isobutyrate (H). Values represent mean ± SD (*n* = 5).

At 24 h, total SCFA production separated treatments into four statistical groups (**Figure 3B**; **Table S1**): Blank and APP alone (group a) < Cellulose (group b) < ternary substrates aPCW and U-aPCW (group c) < Pectin, xyloglucan, and FOS (group d). APP alone produced only minor amounts of SCFAs throughout fermentation (<12.56 mM), and was grouped with the blank in the comparison across all treatments (Table S1). This event indicates minimal fermentation in the absence of carbohydrate substrate. In contrast, FOS rapidly generated high SCFA concentrations by 6 h and remained the strongest producer throughout fermentation (67.45 mM at 24 h), particularly of propionate and butyrate.

Among individual plant cell wall fibres, fermentability reflected substrate composition. Cellulose remained the least fermentable substrate, whereas pectin underwent rapid acetate-dominated fermentation that reached near-maximal SCFA production by 12 h (**Figure 3D**), consistent with its well-described acetogenic profile (Titgemeyer et al., 1991), while producing comparatively lower propionate and butyrate (**Figure 3E, F**). Xyloglucan fermented more gradually but became one of the highest SCFA and butyrate producers after 24 h, in agreement with previous reports (Moro Cantu-Jungles et al., 2019; Sasaki et al., 2025).

Despite sharing identical polysaccharide composition, the two ternary substrates exhibited distinct fermentation kinetics. The hierarchically structured aPCW displayed a gradual, near-linear increase in SCFA production throughout fermentation (aPCW: +0.47 mM·h ¹, r² ≈ 0.99), whereas the unstructured U-aPCW displayed more variable dynamics (+0.13 mM·h ¹, r² = 0.31). Nevertheless, both substrates converged on similar SCFA concentrations by 24 h (49.22 and 47.68 mM, respectively), with comparable acetate and butyrate production (**Figure 3D**, **3F**), indicating that physical assembly influenced fermentation kinetics rather than endpoint metabolite production.

BCFA production displayed the opposite trend to SCFA accumulation **(Figure 3C)**. Cellulose generated the highest concentrations of isobutyrate and isovalerate (2.06 mM total at 24 h; **Figures 3G, H**), whereas highly fermentable substrates (FOS, pectin, and xyloglucan) suppressed BCFA formation relative to the blank. Isobutyrate, however, was gradually increased in xyloglucan (**Figure 3G**), potentially due to xylose fermentation, pointed as a relatively less-fermentable sugar and linked to sustained carbohydrate and protein metabolite production (Zhao et al., 2024). The ternary substrates exhibited intermediate BCFA production.

### 3.3. Fibre composition and physical assembly collectively determine the metabolic response to vacuole-derived (poly)phenols

To quantify the effect of APP, SCFA and BCFA concentrations in APP-treated fermentations were compared with those of the corresponding untreated fermentations, including the Blank, and expressed as absolute differences (ΔAPP, mM; **Figure 4**). Relative changes (log FC) and their significance against untreated fermentations are shown in **Figure S3**. Despite minimal effects on microbial community composition (Section 3.1), APP modulated metabolite production in a substrate-dependent manner, generally promoting saccharolytic fermentation (increased SCFA) while suppressing proteolytic fermentation (decreased BCFA).

**Figure 4.**
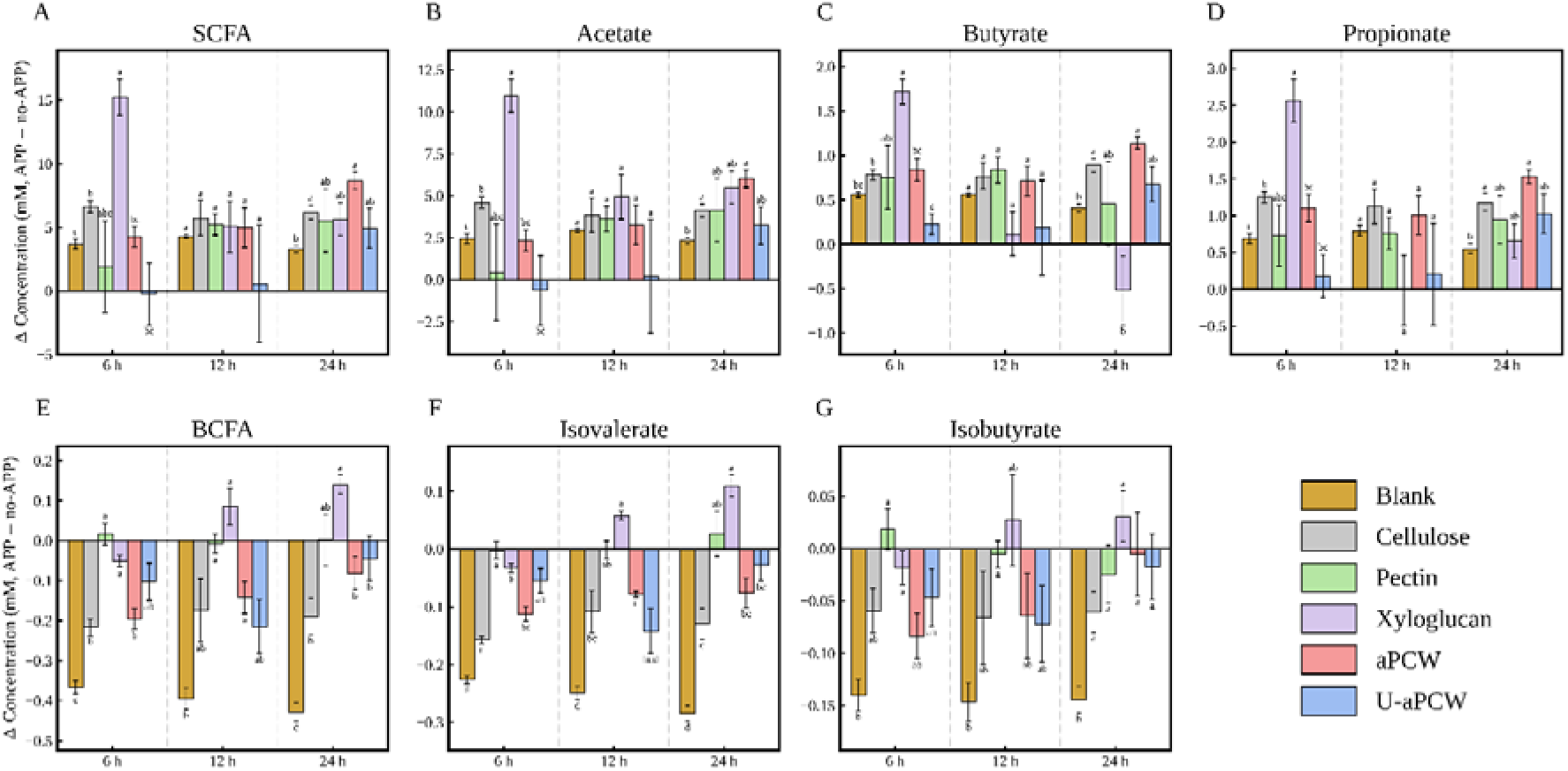
Absolute changes in total and individual SCFA and BCFA concentrations following the addition of vacuole-derived phenolic compounds (APP) across substrates and time points. Values are the difference between the mean concentrations of APP-treated and untreated fermentations (ΔAPP, mM) for the Blank (fermentation without substrate, i.e., the effect of APP alone) and each substrate: total SCFA (A), acetate (B), butyrate (C), propionate (D), total BCFA (E), isovalerate (F) and isobutyrate (G). Total SCFA was calculated as the sum of acetate, propionate and butyrate, and total BCFA as the sum of isobutyrate and isovalerate. Error bars represent the standard error of the difference. Within each time point, bars not sharing a letter differ significantly in the effect of APP, as determined by two-way ANOVA on each pair of conditions (condition × APP interaction) with Holm-adjusted p < 0.05 (*n = 5*).

In the absence of substrate, APP increased total SCFA by 3.3–4.3 mM and reduced total BCFA by 0.37–0.43 mM throughout fermentation (**Figure 4A, E**), the largest relative changes among all conditions (**Figure S3**). Thus, part of the metabolic response to APP was independent of dietary fibre, reflecting either fermentation of extract components or modulation of background metabolism. Against this baseline, APP modulation of metabolite production differed among single-component fibres. On cellulose, APP increased total SCFA production throughout fermentation, exceeding the effect of APP alone at 6 and 24 h (Figure 4A), with significant increases in acetate, propionate, and butyrate (**Figure 4B–D**; **Figure S3**). This indicates that APP “unlocked” the latent SCFA-producing capacity of cellulose fermentation, promoting a shift toward saccharolysis in an otherwise poorly fermentable matrix. In contrast, the response of pectin (+1.9 to +5.5 mM) did not differ from that of APP alone at any time point, with log FC close to 0 (≤ 0.12; **Figure S3**), reflecting a limited scope for improvement on an already rapidly fermented substrate. Xyloglucan displayed the strongest early response, with APP increasing total SCFA production at 6 h (+15.2 mM, about four times the effect of APP alone), largely compensating for its reduced early fermentability.

This effect diminished over time and the effect declined to levels not different from APP alone at 12 and 24 h.

The two ternary substrates again showed different metabolic responses to APP, underscoring the role of physical assembly. In the structured aPCW, the APP effect increased progressively and exceeded that of APP alone at 24 h (+8.7 mM total SCFA, with significant contributions of acetate, propionate and butyrate). The unstructured U-aPCW showed no SCFA response during the first 12 h (−0.2 to +0.6 mM) and a moderate increase at 24 h (+4.9 mM) that did not differ from APP alone.

For BCFA, the pattern was reversed (**Figure 4E–G**). APP alone produced the largest reduction, and the presence of substrate attenuated it at most time points. Reductions remained significant on cellulose (6 and 24 h) and aPCW (6–12 h), statistically different but less intense than with APP alone (Figure 4E; Figure S3). BCFA was essentially unchanged with pectin and increased with xyloglucan at 24 h, mainly through isovalerate. Hence, the reduction in BCFA was largely driven by APP itself, and it weakened as substrate fermentability increased.

### 3.4. Taxa–metabolite correlations define putative saccharolytic and proteolytic guilds in APP-free fermentations substrates

To relate community composition to metabolite output,, we performed taxa–metabolite correlation analysis between filtered taxa and individual metabolites in APP-free fermentations. Correlations were FDR-corrected (**Table S2)** and visualized through hierarchical clustering and network representation (**Figure 5A** and **Figure 5B** respectively). Taxa separated into two main clusters, defined here as putative guilds: Cluster I, comprising taxa positively correlated with SCFA (saccharolytic guild), and Cluster II, comprising taxa negatively correlated with SCFA and/or positively correlated with BCFA (proteolytic guild). Within Cluster I, the strongest SCFA correlates (grouped in subcluster I-A) included *Parabacteroides, Fusicatenibacter, Roseburia, Desulfovibrio,* and *Bacteroides,* each positively associated with all three SCFAs (r > 0.65; p < 0.001). *Lachnospiraceae ND3007 group* and *Blautia* correlated strongly with propionate and butyrate (r > 0.57; p < 0.001), whereas *Faecalibacterium* (acetate-accentuated; r > 0.45; p < 0.05) and *Christensenellaceae R-7* (butyrate-accentuated; r > 0.38; p < 0.01) were also part of this highly saccharolytic group. Subcluster I-B contained weaker but still positive SCFA correlates. For example, within this subcluster *Anaerostipes* showed stronger associations with propionate and butyrate than with acetate, consistent with its ability to utilize acetate during butyrate production (Schwiertz et al., 2002).

**Figure 5.**
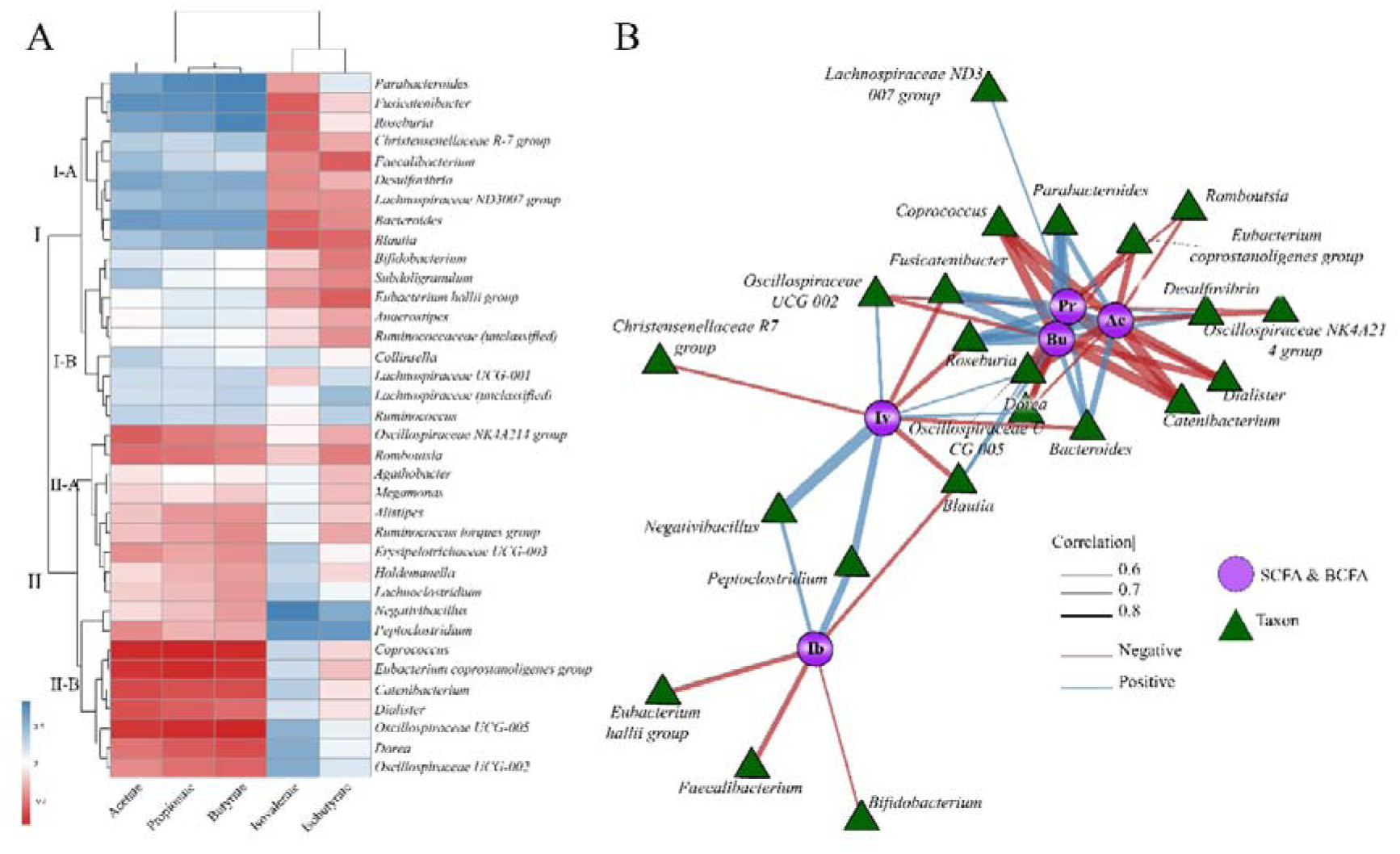
(A) Hierarchical clustering heatmap of Pearson correlation coefficients between log-transformed SCFA and BCFA concentrations and CLR-transformed taxonomic abundances. Zero counts were replaced using the count zero multiplicative (CZM) method prior to CLR transformation. Correlations were calculated across APP-free fermentations at 24 h, and hierarchical clustering was performed using Euclidean distance and complete linkage. The colour scale indicates correlation strength (blue = positive, red = negative). Cluster I (putative saccharolytic guild) and Cluster II (putative proteolytic guild) and their subclusters (I-A, I-B, II-A, II-B) are indicated. (B) Correlation network showing significant associations (FDR-adjusted *p* < 0.05, |*r*| ≥ 0.5). Edge colour indicates correlation direction (blue=positive, red = negative), and edge width is proportional to correlation magnitude. Nodes represent taxa or SCFAs or BCFAs. Network layout was generated using the Fruchterman–Reingold algorithm.

In contrast, within Cluster II, subcluster II-A included taxa with moderate negative SCFA correlations as well as the two taxa most strongly correlated with BCFA, *Negativibacillus* and *Peptoclostridium* (r > 0.59; p < 0.001). Subcluster II-B included taxa mostly negatively associated with SCFA, including *Coprococcus, Eubacterium coprostanoligenes group,* and *Oscillospiraceae UCG-005* (r < −0.81; p < 0.001), as well as taxa positively correlated with isovalerate, such as *Dorea* (r > 0.58; p < 0.01).

Several taxa displayed dual functional patterns. *Fusicatenibacter, Roseburia,* and *Bacteroides* were positively correlated with SCFA while simultaneously negatively correlated with isovalerate (r < −0.54; p < 0.01), indicating a combined promotion of saccharolytic metabolism and suppression of proteolytic byproducts. The strongest significant associations (|r|> 0.50; p < 0.05) are summarized in the network visualization (**Figure 5B**). Hence, rather than being distributed across the microbial community, SCFA and BCFA production was concentrated within discrete groups of taxa rather than distributed across the whole community.

### 3.5. APP induces substrate-dependent shifts in taxa of the saccharolytic and proteolytic guilds

To determine how APP altered fermentation metabolism without extensive community restructuring, differential abundance analysis (DESeq2) was performed for each substrate by comparing fermentations with and without APP. Taxonomic changes were interpreted within the putative fermentation-associated guild framework defined above (**Figure 6, Table S3**). Across 180 log fold-change comparisons, 88.3% ranged between −0.1 and +0.1, with only 14 taxa exhibiting significant differences (*p* < 0.05), consistent with the minimal effects of APP on overall community composition and diversity. While APP alone suppressed *Negativibacillus* and *Peptoclostridium* relative to the Blank (**Figure S2)**, consistent with its reduction of BCFA (**Figure 4E**), no taxon was consistently altered across all substrates. Instead, APP induced coordinated, substrate-dependent changes within the previously identified saccharolytic (Cluster I) and proteolytic (Cluster II) guilds that closely paralleled the observed metabolic responses.

**Figure 6.**
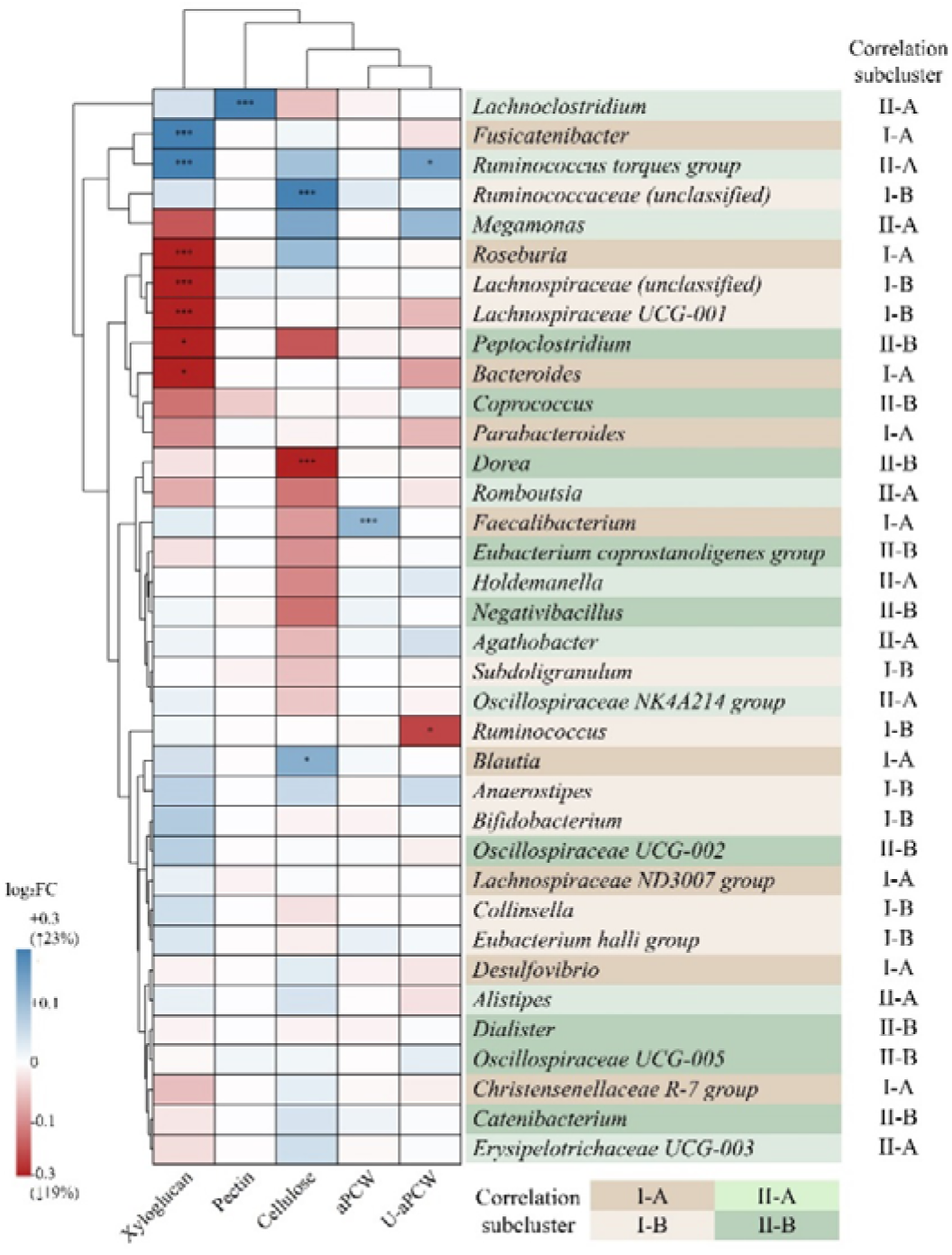
Differential abundance analysis was performed using DESeq2 separately for each fibre substrate, comparing APP-treated versus APP-untreated fermentations. Log fold changes were shrunken using the ashr method. The heatmap displays row-scaled shrunken log fold changes across fibre types, with hierarchical clustering performed using Euclidean distance and complete linkage. Red indicates higher abundance in APP-treated samples, whereas blue indicates higher abundance in APP-untreated samples. Taxon names are colour-coded according to the subclusters identified in Figure 5: brown shades, Cluster I (saccharolytic guild): dark brown, subcluster I-A; light brown, subcluster I-B. Green shades, Cluster II (proteolytic guild): light green, subcluster II-A; dark green, subcluster II-B. Statistical significance is indicated by asterisks (*represents *padj* < 0.05; ** represents padj < 0.01; *** represents *padj* < 0.001; Benjamini–Hochberg adjusted).

This pattern was most evident for cellulose, where APP co-presence produced the strongest metabolic effect (Section 3.3). SCFA-associated taxa from Cluster I generally increased, including significant enrichment of *Blautia* (log FC > 0.19, *p* < 0.05) and unclassified *Ruminococcaceae* (log FC > 0.36, *p* < 0.001), while several protein-associated taxa from Cluster II declined. In particular, *Dorea* decreased significantly (log FC −0.315, *p* < 0.001), accompanied by downward trends in *Peptoclostridium* and *Negativibacillus*, consistent with the simultaneous increase in SCFA production and reduction in BCFA formation. In contrast, pectin—which displayed minimal metabolic responses to APP—showed correspondingly little taxonomic change, with only minor alterations involving low-abundance taxa outside the principal subclusters [I-A and II-A].

Xyloglucan exhibited the largest number of APP-responsive taxa but lacked coordinated regulation at the guild level. Rather than consistently enriching SCFA-associated taxa or suppressing BCFA-associated taxa, APP simultaneously reduced several initially dominant taxa identified in **Figure 2C** and Figure S2 (*Roseburia*, *Parabacteroides*, *Bacteroides*, *Lachnospiraceae* UCG-001 and unclassified *Lachnospiraceae*) while increasing lower-abundance taxa including *Ruminococcus* torques, *Bifidobacterium*, *Collinsella*, *Lachnoclostridium*. This redistribution of community abundance is consistent with the transient metabolic effects observed during xyloglucan fermentation.

The influence of physical fibre assembly became most apparent when comparing the two compositionally matched ternary substrates. Although aPCW and U-aPCW produced similar endpoint communities and metabolite profiles in the absence of APP, APP elicited fundamentally different guild-level responses. In aPCW, APP significantly enriched *Faecalibacterium*, a saccharolytic guild taxon (subcluster I-A) positively correlated with acetate and negatively with isovalerate (**Figure 5**), consistent with the increased SCFA and reduced BCFA production observed during fermentation (**Figure 4)**. In contrast, U-aPCW exhibited only weak metabolic modulation together with a coordinated, although non-significant, reduction across several SCFA-associated taxa, including *Bacteroides*, *Parabacteroides*, *Fusicatenibacter*, and *Desulfovibrio*.

## 4. Discussion

This study showed that: (i) fibre composition establishes the dominant ecological and metabolic trajectory; (ii) physical assembly modulates the pace of fermentation and, critically, determines the efficacy to which APP exerts coordinated functional effects; and (iii) APP acts as a selective ecological modifier, fine-tuning a limited number of taxa associated with fermentation output rather than broadly restructuring communities.. This framework challenges the notion that fibre functions primarily as a facilitator of (poly)phenol metabolism (Cheng et al., 2025; Kilua et al., 2020; Loo et al., 2023; Yang et al., 2021). Instead, it shows that the hierarchical structure of fibre is the primary determinant of microbial and metabolic responses, with polyphenols exerting context-dependent modulation that coincides with shifts in putative saccharolytic and proteolytic guilds. At the poorest end of the fermentability spectrum, cellulose’s rigid and poorly accessible supramolecular structure limited microbial utilization (Holland et al., 2020). This restricted SCFA production and favoured BCFA formation. This metabolic signature, accompanied by the enrichment of proteolytic guild taxa (Cluster II; e.g., *Negativibacillus* and *Peptoclostridium* and *Collinsella*), resembles signatures associated with dysbiotic states and inflammatory bowel disease (J. J. Hou et al., 2024; Hsu et al., 2025; Kim et al., 2024; Pereira et al., 2016). This likely reflects carbohydrate limitation rather than pathogenicity in this closed system. Against this carbohydrate-limited background, APP exerted its strongest effect among the individual fibres. By suppressing Cluster II proteolytic guild taxa, potentially through membrane-disruptive actions as reported for *Peptostreptococcaceae* (Darvishi et al., 2025), while simultaneously increasing that of saccharolytic guild taxa (Cluster I), APP redirected the system toward a more saccharolytic state. This duplibiotic effect (also observed with APP alone), was hence most pronounced when carbohydrate availability was limited. In essence, APP exploited the "ecological space" left by limited nutrient availability or carbohydrate fermentation, rebalancing metabolism through selective modulation of these two opposed guilds.

The duplibiotic pattern of APP was not conserved across all fibre substrates. At the highly fermentable end of the spectrum, represented by pectin, APP exerted only minimal effects. Pectin’s molecular composition presents simpler degradation needs that are met by many dominant gut bacteria orchestrated by polysaccharide utilization loci (PULs) and enzymes (Luis et al., 2017), enabling rapid enzymatic access and acetate-dominated fermentation. This substrate already supported a strongly saccharolytic community with enrichment of saccharolytic guild taxa (Cluster I), while simultaneously suppressing BCFA-correlated *Peptoclostridium*, consistent with previous reports (Tian et al., 2017). Thus, pectin fermentation already achieved what APP would otherwise promote, a high SCFA production and low BCFA, leaving little ecological room for additional modulation. This ceiling effect explains why APP contributed relatively little to pectin fermentation. Previous work suggests that substantially higher pectin-to-(poly)phenol ratios, at least 1:1, may be required to detect measurable synergy (J. Wu et al., 2025). In other rapidly fermentable substrates such as inulin, (poly)phenol antimicrobial activity may even suppress fermentation (Nagata et al., 2018), possibly reflecting competition between saccharolytic expansion and phenolic inhibition.

Xyloglucan occupied a middle position on the fermentability spectrum, reflecting the need for greater metabolic and microbial specialization for its degradation. Its xylose-substituted glucan backbone requires coordinated debranching and backbone hydrolysis, involving xyloglucanases and accessory enzymes such as xylanases and xylosidases (Déjean et al., 2019; Kaoutari et al., 2013; Moro Cantu-Jungles et al., 2019). This results in slower but ultimately efficient fermentation through both primary degradation and cross-feeding, which could support SCFA delivery to distal colonic regions. APP produced its strongest early metabolic effect with this substrate, substantially promoting SCFA and suppressing BCFA at 6 h, but these effects were transient and diminished by 24 h. The taxonomic basis for this early shift remains unclear, as 16S profiling was only performed at the endpoint. At 24 h, APP exhibited redistributed abundance among dominant and less abundant taxa without a consistent guild-level direction, a pattern more consistent with ecological buffering (Steinert et al., 2025), than with directional shifts in either guild. This transient pattern further supports the inverse relationship between fermentability and APP efficacy.

Beyond molecular composition, a second layer of structural control emerged. Despite identical polysaccharide composition, only the structurally assembled aPCW enabled a consistent APP response in both metabolite and taxa modulation, while the unstructured U-aPCW remained largely unresponsive. Because APP was already intestinally bioaccessible, simple (poly)phenol–fibre entrapment is unlikely to explain this difference. Instead, we first demonstrated how physical assembly regulates the tempo of microbial substrate access. The more linear fermentation kinetics of aPCW suggest that the assembled cell-wall network enabled more coordinated and sustained exposure of fermentable domains, whereas U-aPCW, lacking this higher-order structure, provided more immediate but less structured substrate access. In line with this, it has been recognized that intact structures are fermented slower than disrupted structures (Dagbasi et al., 2020, 2024; Lu et al., 2020; Mikkelsen et al., 2011) and that this structure further creates positive ecological niches for gut bacterial growth (Bulut et al., 2023). Thus, the temporal coordination imposed by physical assembly appears to be a critical factor enabling APP to exert guild-level modulation.

Our results contrast with a study where polyphenols exerted minimal microbial modulation on isolated apple cell walls, despite showing a positive effect on cellulose (Phan et al., 2020). This discrepancy likely reflects that isolation processing disrupted the hierarchical structure of the cell walls, analogous to the unresponsive U-aPCW in our study. Similarly, divergent responses have been reported for soluble versus insoluble fibre mixtures supplemented with phenolic acids (Liu et al., 2025), though the basis for this differential effect remains unclear. Together, these observations underscore the need to consider physical assembly—rather than composition alone—when evaluating fibre–(poly)phenol interactions, a dimension that remains minimally explored in the literature (Cheng et al., 2025). Our direct comparison of matched structured and unstructured substrates provides, to our knowledge, the first controlled evidence that physical assembly determines whether (poly)phenols can exert coordinated functional effects.

It should be highlighted however that our batch in vitro system from a three-donor poool does not permit direct inference of host outcomes, a limitation shared with most reductionist fermentation studies. Consequently, inter-individual variability in fibre could not be elucidated, and the substrate-dependent responses reported here should be regarded as specific to this pooled community. Further research across independent human microbiomes (e.g., individual donors, or pools from contrasting microbiomes) is warranted to confirm the reproducibility of these responses. Moreover, SCFA and BCFA were the only fermentation products quantified; BCFA provide only a partial indication of protein fermentation, and other markers (e.g., ammonia, phenolic and indolic metabolites) are suggested to emplo. In addition, guild membership was inferred from correlations between 16S rRNA gene profiles and metabolites across substrates, which may partly reflect shared responses to substrate rather than direct metabolic contributions.

Nevertheless, as discussed above, these observations are consistent with previous studies and with the emerging view that phenolic compounds act as subtle ecological modulators with both trophic and antimicrobial activities, rather than primary drivers of community structure, within microbiotas in which a limited number of guilds are associated with community metabolism and host-associated phenotypes (Wang et al., 2026; G. Wu et al., 2024; H. Zhang et al., 2026), and help resolve open questions in the literature, challenging the notion that fibre functions primarily as a facilitator of (poly)phenol metabolism. Here, strong associations with SCFA and BCFA were concentrated in a limited number of taxa, and APP-induced shifts in these taxa paralleled changes in metabolite output. This supports a guild-based approach to interpreting fibre–(poly)phenol interactions: first identify metabolite-associated guilds, then interpret metabolic effects through shifts within them, since relatively small shifts may alter fermentation output without large-scale community restructuring. Importantly, the same (poly)phenol exposure produced different outcomes depending on whether the carbohydrate environment was rapidly fermentable, structurally constrained or physically assembled, implying that neither the effects of dietary (poly)phenols nor those of dietary fibre can be interpreted in isolation.

## 5. Conclusions

This study supports a hierarchical framework in which dietary fibre composition and physical assembly shape colonic fermentation and condition microbial responses to apple (poly)phenols. In human faecal batch fermentations, APP altered microbial metabolic outputs despite having limited effects on overall community composition and diversity. Increases in SCFA and decreases in BCFA concentrations were accompanied by changes in the relative abundance of taxa assigned to putative saccharolytic and proteolytic guilds based on taxon–metabolite correlations. These responses depended on the polysaccharide system, being most pronounced during cellulose fermentation and transient with xyloglucan.

A key finding was the response of the compositionally matched plant cell wall systems: coordinated metabolic and taxonomic changes were observed with the physically assembled analogue (aPCW), whereas its unstructured counterpart showed limited responses. These findings support a role for higher order organisation in determining both fermentation kinetics and microbial responses to co-supplied (poly)phenols. They provide a basis for considering physical assembly alongside polysaccharide composition when designing fibre–(poly)phenol combinations to modulate microbial metabolism. Validation across individual donor microbiomes and *in vivo* is needed to establish the generalisability and nutritional relevance of these effects.

## CRediT authorship contribution statement

**Ivan M. Lopez-Rodulfo**: Investigation, Methodology, Visualization, Writing – original draft. **Clay Swackhamer**: Investigation, Methodology, Visualization. **Julia Bechtner:** Methodology. **Fang Fang:** Methodology. **Bruce Hamaker**: Methodology, Supervision, Writing – review & editing. **Mario M. Martinez**: Conceptualization, Methodology, Supervision, Visualization, Funding acquisition, Writing – review & editing.

## Data availability statement

All data needed to evaluate the conclusions of this paper are presented in the paper and/or the Supplementary Materials. Raw reads are available in the National Center for Biotechnology Information Sequence Read Archive (NCBI; SRA), BioProject PRJNA1506331, and BioSamples SAMN62124126 to SAMN62124204. These data are available at: http://www.ncbi.nlm.nih.gov/bioproject/1506331. Metabolite datasets and R scripts used for statistical analysis are available from the corresponding author upon reasonable request.

## References

Aprikian, O., Duclos, V., Guyot, S., Besson, C., Manach, C., Bernalier, A., Morand, C., Rémésy, C., & Demigné, C. (2003). Apple pectin and a polyphenol-rich apple concentrate are more effective together than separately on cecal fermentations and plasma lipids in rats. Journal of Nutrition, 133(6), 1860–1865. 10.1093/jn/133.6.1860

Bechtner, J. D., Lopez-Rodulfo, I. M., Gallego-Lobillo, P., Hosek, J., Schwab, C., & Martinez, M. M. (2025). Polyphenols non-covalently bound to artificial plant cell walls are readily accessible to fecal microbiota and transiently alter fermentation activity. Food Chemistry, 493(P2), 145817. 10.1016/j.foodchem.2025.145817

Bokulich, N. A., Kaehler, B. D., Rideout, J. R., Dillon, M., Bolyen, E., Knight, R., Huttley, G. A., & Gregory Caporaso, J. (2018). Optimizing taxonomic classification of marker-gene amplicon sequences with QIIME 2’s q2-feature-classifier plugin. Microbiome, 6(1), 1–17. 10.1186/s40168-018-0470-z

Bulut, N., Cantu-Jungles, T. M., Zhang, X., Mutlu, Z., Cakmak, M., & Hamaker, B. R. (2023). Matrix-entrapped fibers create ecological niches for gut bacterial growth. Scientific Reports, 13(1), 1–12. 10.1038/s41598-023-27907-7

Callahan, B. J., McMurdie, P. J., Rosen, M. J., Han, A. W., Johnson, A. J. A., & Holmes, S. P. (2016). DADA2: High-resolution sample inference from Illumina amplicon data. Nature Methods, 13(7), 581–583. 10.1038/nmeth.3869

Cantu-Jungles, T. M., & Hamaker, B. R. (2020). New View on Dietary Fiber Selection for Predictable Shifts in Gut Microbiota. MBio, 11(1). 10.1128/mBio.02179-19

Cantu-Jungles, T. M., & Hamaker, B. R. (2023). Tuning Expectations to Reality: Don’t Expect Increased Gut Microbiota Diversity with Dietary Fiber. The Journal of Nutrition, 153(11), 3156–3163. 10.1016/j.tjnut.2023.09.001

Cheng, B., Feng, H., Li, C., Jia, F., & Zhang, X. (2025). The mutual effect of dietary fiber and polyphenol on gut microbiota: Implications for the metabolic and microbial modulation and associated health benefits. Carbohydrate Polymers, 358(January), 123541. 10.1016/j.carbpol.2025.123541

Dagbasi, A., Byrne, C., Blunt, D., Serrano-Contreras, J. I., Becker, G. F., Blanco, J. M., Camuzeaux, S., Chambers, E., Danckert, N., Edwards, C., Bernal, A., Garcia, M. V., Hanyaloglu, A., Holmes, E., Ma, Y., Marchesi, J., Martinez-Gili, L., Mendoza, L., Tashkova, M.,… Frost, G. (2024). Diet shapes the metabolite profile in the intact human ileum, which affects PYY release. Science Translational Medicine, 16(752), 1–16. 10.1126/scitranslmed.adm8132

Dagbasi, A., Lett, A. M., Murphy, K., & Frost, G. (2020). Understanding the interplay between food structure, intestinal bacterial fermentation and appetite control. Proceedings of the Nutrition Society, 79(4), 514–530. 10.1017/S0029665120006941

Darvishi, M., Rafsanjani, S. M. R. H., Nouri, M., Abbaszadeh, S., Heidari-Soureshjani, S., Kasiri, K., & Rahimian, G. (2025). Biological Mechanisms of Polyphenols against Clostridium Difficile: A Systematic Review. Infectious Disorders - Drug Targets, 25(3). 10.2174/0118715265313944240726115600

Deehan, E. C., Yang, C., Perez-Muñoz, M. E., Nguyen, N. K., Cheng, C. C., Triador, L., Zhang, Z., Bakal, J. A., & Walter, J. (2020). Precision Microbiome Modulation with Discrete Dietary Fiber Structures Directs Short-Chain Fatty Acid Production. Cell Host & Microbe, 27(3), 389–404. 10.1016/j.chom.2020.01.006

Déjean, G., Tauzin, A. S., Bennett, S. W., Creagh, A. L., & Brumer, H. (2019). Adaptation of syntenic xyloglucan utilization loci of human gut Bacteroidetes to polysaccharide side chain diversity. Applied and Environmental Microbiology, 85(20), 1–17. 10.1128/AEM.01491-19

Diener, M., Adamcik, J., Sánchez-Ferrer, A., Jaedig, F., Schefer, L., & Mezzenga, R. (2019). Primary, Secondary, Tertiary and Quaternary Structure Levels in Linear Polysaccharides: From Random Coil, to Single Helix to Supramolecular Assembly. Biomacromolecules, 20(4), 1731–1739. 10.1021/acs.biomac.9b00087

Elshahed, M. S., Miron, A., Clara, A., & Farag, M. A. (2021). Pectin in diet : Interactions with the human microbiome, role in gut homeostasis, and nutrient-drug interactions. Carbohydrate Polymers, 255(November 2020), 117388. 10.1016/j.carbpol.2020.117388

Fang, F., Franco, M., Gallego-Lobillo, P., Spotti, M. J., Bechtner, J. D., & Martinez, M. M. (2026). Interactive effects of free and cell wall-bound polyphenols on starch pasting, retrogradation, gelation and digestibility. Food Hydrocolloids, 175(October 2025), 112457. 10.1016/j.foodhyd.2026.112457

Fu, J., Zheng, Y., Gao, Y., & Xu, W. (2022). Dietary Fiber Intake and Gut Microbiota in Human Health. Microorganisms, 10(12), 2507. 10.3390/microorganisms10122507

García-Cordero, J., Mateos, R., González-Rámila, S., Seguido, M. A., Sierra-Cinos, J. L., Sarriá, B., & Bravo, L. (2023). Dietary Supplements Containing Oat Beta-Glucan and/or Green Coffee (Poly)phenols Showed Limited Effect in Modulating Cardiometabolic Risk Biomarkers in Overweight/Obese Patients without a Lifestyle Intervention. Nutrients, 15(9), 2223. 10.3390/nu15092223

Gidley, M. J., & Yakubov, G. E. (2019). Functional categorisation of dietary fibre in foods: Beyond ‘soluble’ vs ‘insoluble.’ Trends in Food Science & Technology, 86, 563–568. 10.1016/j.tifs.2018.12.006

Hamaker, B. R., & Tuncil, Y. E. (2014). A perspective on the complexity of dietary fiber structures and their potential effect on the gut microbiota. In Journal of Molecular Biology (Vol. 426, Number 23, pp. 3838–3850). Academic Press. 10.1016/j.jmb.2014.07.028

Holland, C., Ryden, P., Edwards, C. H., & Grundy, M. M. L. (2020). Plant cell walls: Impact on nutrient bioaccessibility and digestibility. Foods, 9(2), 1–16. 10.3390/foods9020201

Hou, C., Shi, H., Xiao, J., Song, X., Luo, Z., Ma, X., Shi, L., Wei, H., & Li, J. (2024). Pomegranate Juice Supplemented with Inulin Modulates Gut Microbiota and Promotes the Production of Microbiota-Associated Metabolites in Overweight/Obese Individuals: A Randomized, Double-Blind, Placebo-Controlled Trial. Journal of Agricultural and Food Chemistry, 72(26), 14663–14677. 10.1021/acs.jafc.4c00849

Hou, J. J., Ding, L., Yang, T., Yang, Y. F., Jin, Y. P., Zhang, X. P., Ma, A. H., & Qin, Y. H. (2024). The proteolytic activity in inflammatory bowel disease: insight from gut microbiota. Microbial Pathogenesis, 188(January), 106560. 10.1016/j.micpath.2024.106560

Hsu, B. M., Chen, J. S., Wang, W. Y., Chen, C. J., Fan, C. W., Wu, C. C., Hussain, B., & Tsai, H. C. (2025). An integral view of gut microbiome diversity and functional metabolic changes of a gut-brain axis associated with dementia based on metagenomic analysis. Physiology and Behavior, 302(July), 115112. 10.1016/j.physbeh.2025.115112

Johnson, K. L., Gidley, M. J., Bacic, A., & Doblin, M. S. (2018). Cell wall biomechanics: a tractable challenge in manipulating plant cell walls ‘fit for purpose’! Current Opinion in Biotechnology, 49, 163–171. 10.1016/j.copbio.2017.08.013

Kaoutari, A. El, Armougom, F., Gordon, J. I., Raoult, D., & Henrissat, B. (2013). The abundance and variety of carbohydrate-active enzymes in the human gut microbiota. Nature Reviews Microbiology, 11(7), 497–504. 10.1038/nrmicro3050

Karl, J. P., Fagnant, H. S., Radcliffe, P. N., Wilson, M., Karis, A. J., Sayers, B., Wijeyesekera, A., Gibson, G. R., Lieberman, H. R., Giles, G. E., & Soares, J. W. (2025). Gut microbiota-targeted dietary supplementation with fermentable fibers and polyphenols prevents hypobaric hypoxia-induced increases in intestinal permeability. American Journal of Physiology-Regulatory, Integrative and Comparative Physiology, 329(3), R378–R399. 10.1152/ajpregu.00109.2025

Kaur, A., Chen, T., Green, S. J., Mutlu, E., Martin, B. R., Rumpagaporn, P., Patterson, J. A., Keshavarzian, A., & Hamaker, B. R. (2019). Physical Inaccessibility of a Resistant Starch Shifts Mouse Gut Microbiota to Butyrogenic Firmicutes. Molecular Nutrition & Food Research, 63(7). 10.1002/mnfr.201801012

Kilua, A., Han, K., & Fukushima, M. (2020). Effect of polyphenols isolated from purple sweet potato ( Ipomoea batatas cv. Ayamurasaki) on the microbiota and the biomarker of colonic fermentation in rats fed with cellulose or inulin. Food & Function, 11(11), 10182–10192. 10.1039/D0FO02111C

Kim, H. S., Oh, S. J., Kim, B. K., Kim, J. E., Kim, B., Park, Y., Yang, B., Lee, J.-Y., Bae, J., & Lee, C. K. (2024). Dysbiotic signatures and diagnostic potential of gut microbial markers for inflammatory bowel disease in Korean population. Scientific Reports, 14(1), 23701. 10.1038/s41598-024-74002-6

Liu, S., Perez Donado, C. E., & Rose, D. J. (2025). Impact of ferulic and vanillic acids on soluble and insoluble dietary fiber utilization from maize bran by the human gut microbiota. Food and Function, 16(13), 5409–5420. 10.1039/d5fo00005j

Loo, Y. T., Howell, K., Suleria, H., Zhang, P., Liu, S., & Ng, K. (2023). Fibre fermentation and pig faecal microbiota composition are affected by the interaction between sugarcane fibre and (poly)phenols in vitro. International Journal of Food Sciences and Nutrition, 74(2), 219–233. 10.1080/09637486.2023.2187329

Lopez-Rodulfo, I. M., Rasmussen, M. K., Hamaker, B., & Martinez, M. M. (2026). Selective detoxification of digesta revealed how cold-pressed apple fractions modulate transepithelial glucose transport and stimulate GLP-1 secretion. Food Chemistry, 509, 148588. 10.1016/j.foodchem.2026.148588

Lopez-Rodulfo, I. M., Stentoft, E. W., & Martinez, M. M. (2025). Comparative assessment of polyphenol bioaccessibility in cold-pressed apple fractions using static and semi-dynamic digestion models. Food Research International, 202(January), 115743. 10.1016/j.foodres.2025.115743

Lopez-Rodulfo, I. M., Tsochatzis, E. D., Stentoft, E. W., Martinez-Carrasco, P., Bechtner, J. D., & Martinez, M. M. (2024). Partitioning and in vitro bioaccessibility of apple polyphenols during mechanical and physiological extraction: A hierarchical clustering analysis with LC-ESI-QTOF-MS/MS. Food Chemistry, 441(January), 138320. 10.1016/j.foodchem.2023.138320

Lopez-Siles, M., Khan, T. M., Duncan, S. H., Harmsen, H. J. M., Garcia-Gil, L. J., & Flint, H. J. (2012). Cultured Representatives of Two Major Phylogroups of Human Colonic Faecalibacterium prausnitzii Can Utilize Pectin, Uronic Acids, and Host-Derived Substrates for Growth. Applied and Environmental Microbiology, 78(2), 420–428. 10.1128/AEM.06858-11

Love, M. I., Huber, W., & Anders, S. (2014). Moderated estimation of fold change and dispersion for RNA-seq data with DESeq2. Genome Biology, 15(12), 550. 10.1186/s13059-014-0550-8

Lu, S., Flanagan, B. M., Williams, B. A., Mikkelsen, D., & Gidley, M. J. (2020). Cell wall architecture as well as chemical composition determines fermentation of wheat cell walls by a faecal inoculum. Food Hydrocolloids, 107(March), 105858. 10.1016/j.foodhyd.2020.105858

Luis, A. S., Briggs, J., Zhang, X., Farnell, B., Ndeh, D., Labourel, A., Baslé, A., Cartmell, A., Terrapon, N., Stott, K., Lowe, E. C., McLean, R., Shearer, K., Schückel, J., Venditto, I., Ralet, M.-C., Henrissat, B., Martens, E. C., Mosimann, S. C.,… Gilbert, H. J. (2017). Dietary pectic glycans are degraded by coordinated enzyme pathways in human colonic Bacteroides. Nature Microbiology, 3(2), 210–219. 10.1038/s41564-017-0079-1

McDonald, D., Price, M. N., Goodrich, J., Nawrocki, E. P., Desantis, T. Z., Probst, A., Andersen, G. L., Knight, R., & Hugenholtz, P. (2012). An improved Greengenes taxonomy with explicit ranks for ecological and evolutionary analyses of bacteria and archaea. ISME Journal, 6(3), 610–618. 10.1038/ismej.2011.139

Mikkelsen, D., Gidley, M. J., & Williams, B. A. (2011). In Vitro Fermentation of Bacterial Cellulose Composites as Model Dietary Fibers. Journal of Agricultural and Food Chemistry, 59(8), 4025–4032. 10.1021/jf104855e

Mikkelsen, D., Lopez-Sanchez, P., Wang, D., & Gidley, M. J. (2020). Formation of Cellulose-Based Composites with Hemicelluloses and Pectins Using Komagataeibacter Fermentation (pp. 73–87). 10.1007/978-1-0716-0621-6_5

Moro Cantu-Jungles, T., do Nascimento, G. E., Zhang, X., Iacomini, M., Cordeiro, L. M. C., & Hamaker, B. R. (2019). Soluble xyloglucan generates bigger bacterial community shifts than pectic polymers during in vitro fecal fermentation. Carbohydrate Polymers, 206(October 2018), 389–395. 10.1016/j.carbpol.2018.11.011

Nagata, R., Echizen, M., Yamaguchi, Y., Han, K. H., Shimada, K., Ohba, K., Kitano-Okada, T., Nagura, T., Uchino, H., & Fukushima, M. (2018). Effect of a combination of inulin and polyphenol-containing adzuki bean extract on intestinal fermentation in vitro and in vivo. Bioscience, Biotechnology and Biochemistry, 82(3), 489–496. 10.1080/09168451.2018.1429886

Padayachee, A., Day, L., Howell, K., & Gidley, M. J. (2017). Complexity and health functionality of plant cell wall fibers from fruits and vegetables. Critical Reviews in Food Science and Nutrition, 57(1), 59–81. 10.1080/10408398.2013.850652

Pereira, F. L., Oliveira Júnior, C. A., Silva, R. O. S., Dorella, F. A., Carvalho, A. F., Almeida, G. M. F., Leal, C. A. G., Lobato, F. C. F., & Figueiredo, H. C. P. (2016). Complete genome sequence of Peptoclostridium difficile strain Z31. Gut Pathogens, 8(1), 11. 10.1186/s13099-016-0095-3

Phan, A. D. T., Williams, B. A., Netzel, G., Mikkelsen, D., D’Arcy, B. R., & Gidley, M. J. (2020). Independent fermentation and metabolism of dietary polyphenols associated with a plant cell wall model. Food and Function, 11(3), 2218–2230. 10.1039/c9fo02987g

Pihelgas, S., Ehala-Aleksejev, K., Adamberg, S., Kazantseva, J., & Adamberg, K. (2024). The gut microbiota of healthy individuals remains resilient in response to the consumption of various dietary fibers. Scientific Reports, 14(1), 22208. 10.1038/s41598-024-72673-9

Reichardt, N., Vollmer, M., Holtrop, G., Farquharson, F. M., Wefers, D., Bunzel, M., Duncan, S. H., Drew, J. E., Williams, L. M., Milligan, G., Preston, T., Morrison, D., Flint, H. J., & Louis, P. (2018). Specific substrate-driven changes in human faecal microbiota composition contrast with functional redundancy in short-chain fatty acid production. The ISME Journal, 12(2), 610–622. 10.1038/ismej.2017.196

Renard, C. M. G. ., Baron, A., Guyot, S., & Drilleau, J.-F. (2001). Interactions between apple cell walls and native apple polyphenols: quantification and some consequences. International Journal of Biological Macromolecules, 29(2), 115–125. 10.1016/S0141-8130(01)00155-6

Rivas-Santisteban, J., Yubero, P., Robaina-Estévez, S., González, J. M., Tamames, J., & Pedrós-Alió, C. (2024). Quantifying microbial guilds. ISME Communications, 4(1). 10.1093/ismeco/ycae042

Rodríguez-Daza, M. C., Pulido-Mateos, E. C., Lupien-Meilleur, J., Guyonnet, D., Desjardins, Y., & Roy, D. (2021). Polyphenol-Mediated Gut Microbiota Modulation: Toward Prebiotics and Further. Frontiers in Nutrition, 8(June). 10.3389/fnut.2021.689456

Rose, D. J., Patterson, J. A., & Hamaker, B. R. (2010). Structural Differences among Alkali-Soluble Arabinoxylans from Maize ( Zea mays ), Rice ( Oryza sativa ), and Wheat ( Triticum aestivum ) Brans Influence Human Fecal Fermentation Profiles. Journal of Agricultural and Food Chemistry, 58(1), 493–499. 10.1021/jf9020416

Sasaki, H., Masutomi, H., Kobayashi, Y., Toyohara, K., Imaizumi, K., Nakayama, Y., & Ishihara, K. (2025). Evaluation of the Fermentation Characteristics of Prebiotic-Containing Granola and Short-Chain Fatty Acid Production in an In Vitro Gut Microbiota Model. Food Science and Nutrition, 13(5). 10.1002/fsn3.70252

Schwiertz, A., Hold, G. L., Duncan, S. H., Gruhl, B., Collins, M. D., Lawson, P. A., Flint, H. J., & Blaut, M. (2002). Anaerostipes caccae gen. nov., sp. nov., a New Saccharolytic, Acetate-utilising, Butyrate-producing Bacterium from Human Faeces. Systematic and Applied Microbiology, 25(1), 46–51. 10.1078/0723-2020-00096

Solvang, M., Farquharson, F. M., Horgan, G., Pisano, S., Holck, J., Zeuner, B., Russell, W. R., & Louis, P. (2025). Roles of human colonic bacteria in pectin utilization and associated cross-feeding networks revealed using synthetic co-cultures. Microbiology, 171(5), 1–16. 10.1099/mic.0.001559

Steinert, R. E., Rehman, A., Sadabad, M. S., Milanese, A., Wittwer-Schegg, J., Burton, J. P., & Spooren, A. (2025). Microbial micronutrient sharing, gut redox balance and keystone taxa as a basis for a new perspective to solutions targeting health from the gut. Gut Microbes, 17(1). 10.1080/19490976.2025.2477816

Tian, L., Bruggeman, G., van den Berg, M., Borewicz, K., Scheurink, A. J. W., Bruininx, E., de Vos, P., Smidt, H., Schols, H. A., & Gruppen, H. (2017). Effects of pectin on fermentation characteristics, carbohydrate utilization, and microbial community composition in the gastrointestinal tract of weaning pigs. Molecular Nutrition and Food Research, 61(1), 1–10. 10.1002/mnfr.201600186

Titgemeyer, E., Bourquin, L., Fahey, G., & Garleb, K. (1991). Fermentability of various fiber sources by human fecal bacteria in vitro. The American Journal of Clinical Nutrition, 53(6), 1418–1424. 10.1093/ajcn/53.6.1418

Tuncil, Y. E., Nakatsu, C. H., Kazem, A. E., Arioglu-Tuncil, S., Reuhs, B., Martens, E. C., & Hamaker, B. R. (2017). Delayed utilization of some fast-fermenting soluble dietary fibers by human gut microbiota when presented in a mixture. Journal of Functional Foods, 32, 347–357. 10.1016/j.jff.2017.03.001

von Muhlenbrock, C., Aronsohn, F., Quera, R., & Madrid, A. M. (2025). The role of dietary fiber in the gastrointestinal tract: when, how and why? Best Practice & Research Clinical Gastroenterology, 79, 102080. 10.1016/j.bpg.2025.102080

Walsh, S. K., Armet, A. M., Nikolaeva, D. D., Mota, J. F., Lucey, A. J., Oliero, M., & Walter, J. (2026). Optimizing Dietary Fiber Intake: Strategies for Human Nutrition and Food Science. Annual Review of Food Science and Technology, 17(1), 25–50. 10.1146/annurev-food-052824-044842

Wang, T., Gyori, B., Weiss, S. T., Menichetti, G., & Liu, Y.-Y. (2026). Revealing interactions between microbes, metabolites, and dietary compounds using genome-scale analysis. Microbiome, 14(1), 74. 10.1186/s40168-025-02312-6

Wu, G., Xu, T., Zhao, N., Lam, Y. Y., Ding, X., Wei, D., Fan, J., Shi, Y., Li, X., Li, M., Ji, S., Wang, X., Fu, H., Zhang, F., Shi, Y., Zhang, C., Peng, Y., & Zhao, L. (2024). A core microbiome signature as an indicator of health. Cell, 187(23), 6550–6565.e11. 10.1016/j.cell.2024.09.019

Wu, J., Shen, S., Cheng, H., Pan, H., & Ye, X. (2025). RG-I pectic polysaccharides and hesperidin synergistically modulate gut microbiota : An in vitro study targeting the proportional relationship. Food Chemistry, 462(August 2024), 141010. 10.1016/j.foodchem.2024.141010

Yang, Z., Huang, T., Li, P., Ai, J., Liu, J., Bai, W., & Tian, L. (2021). Dietary Fiber Modulates the Fermentation Patterns of Cyanidin-3-O-Glucoside in a Fiber-Type Dependent Manner. Foods, 10(6), 1386. 10.3390/foods10061386

Zduńczyk, Z., Juśkiewicz, J., & Estrella, I. (2006). Cecal parameters of rats fed diets containing grapefruit polyphenols and inulin as single supplements or in a combination. Nutrition, 22(9), 898–904. 10.1016/j.nut.2006.05.010

Zhang, H., Yan, X., Hou, Y., Ou, C., Luo, S., Liu, C., Li, Z., Zhang, G., & Chen, T. (2026). Polyphenols Nonmonotonically Modulate the Inulin-Driven Gut Microbial Network In Vivo. Journal of Agricultural and Food Chemistry, 74(9), 7530–7542. 10.1021/acs.jafc.5c13003

Zhang, W., Luo, H., Keung, W., Chan, Y., Chan, K., Xiao, X., Li, F., Lyu, A., Dong, C., & Xu, J. (2025). Impact of pectin structural diversity on gut microbiota : A mechanistic exploration through in vitro fermentation. Carbohydrate Polymers, 355(January), 123367. 10.1016/j.carbpol.2025.123367

Zhang, X., Dong, L., Jia, X., Liu, L., Chi, J., Huang, F., Ma, Q., Zhang, M., & Zhang, R. (2020). Bound Phenolics Ensure the Antihyperglycemic Effect of Rice Bran Dietary Fiber in db / db Mice via Activating the Insulin Signaling Pathway in Skeletal Muscle and Altering Gut Microbiota. Journal of Agricultural and Food Chemistry, 68(15), 4387–4398. 10.1021/acs.jafc.0c00584

Zhao, Y., Wang, Y., Ma, Q., Wang, D., Jiang, Q., Wang, P., Ge, Z., Wang, J., Qin, P., & Zhao, X. (2024). Different microbiota modulation and metabolites generation of five dietary glycans during in vitro gut fermentation are determined by their monosaccharide profiles. Food Research International, 196(50), 115011. 10.1016/j.foodres.2024.115011

