## Supplementary information for "Dietary fibre hierarchical structure determines gut microbiota responses to apple (poly)phenols through metabolite-associated microbial guilds"

This PDF file includes:

I. Supplementary Figures and Legends

Supplementary Figure 1. Bacillota/Bacteroidota ratio (formerly Firmicutes/Bacteroidota).

Supplementary Figure 2. Differential abundance of gut bacterial taxa in response to dietary fiber treatments.

II. Supplementary Tables and Legends

Supplementary Table 1. Concentrations (mM) of short-chain fatty acids (SCFA) and branched-chain fatty acids (BCFA).

Supplementary Table 2. Pearson correlation coefficients (r) and corresponding p-values for associations between gut microbial taxa and individual microbial metabolites (SCFAs and BCFAs).

Supplementary Table 3. Shrunken log₂ fold changes of microbial taxa in APP-treated versus untreated fermentations across different fiber substrates (DESeq2 analysis).

Figure S1. Bacillota/Bacteroidota ratio (formerly Firmicutes/Bacteroidota). Statistical differences among treatments were assessed using Tukey’s multiple comparisons test. Bars not sharing the same letter differ significantly (p < 0.05).

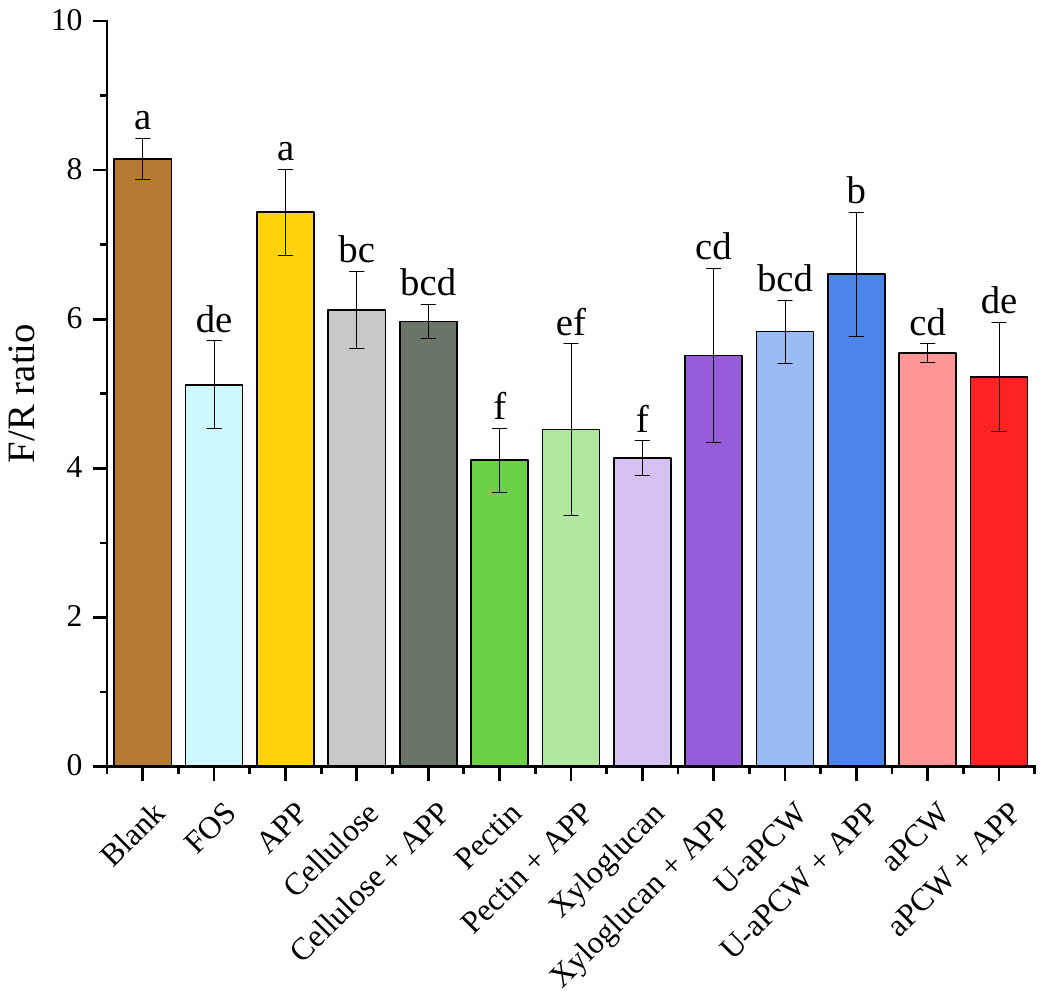

Figure S2. Differential abundance of gut bacterial taxa in response to dietary fiber treatments. Bar plots show log2 fold change (vs. blank control) for each fiber type. Only samples without APP are included. Taxa are ordered by effect size

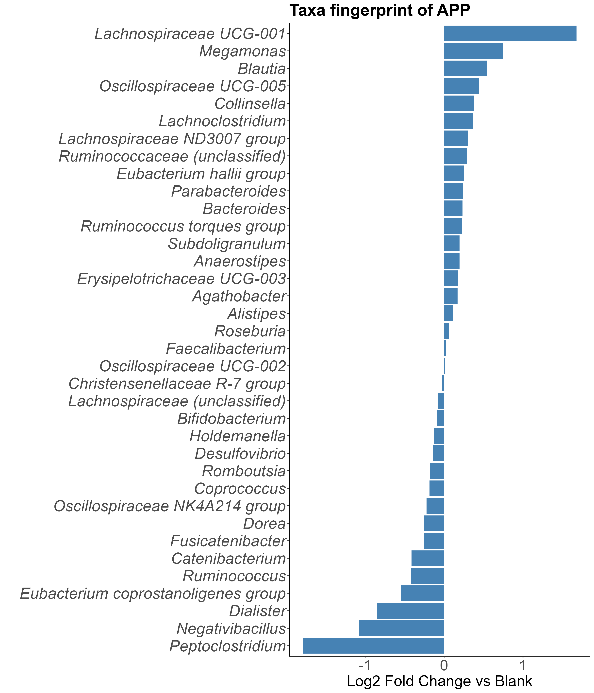

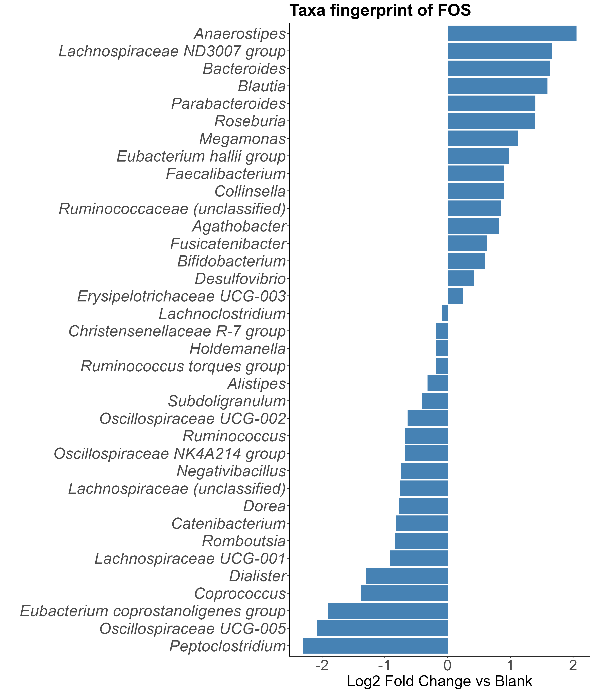

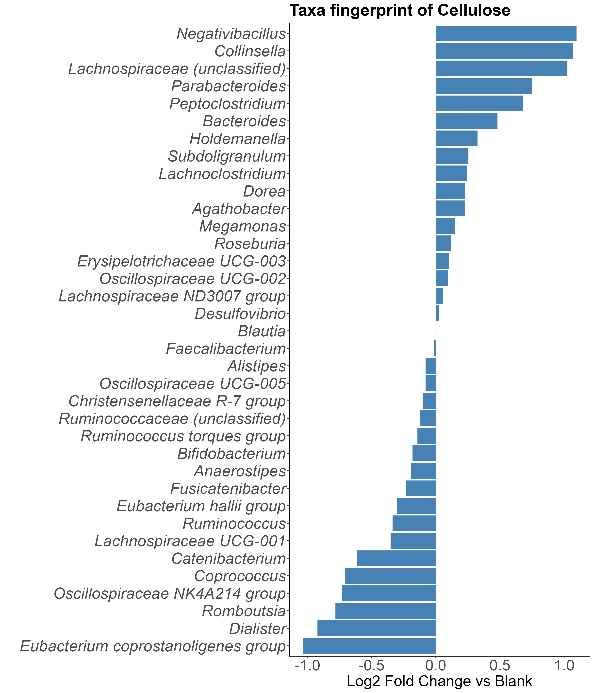

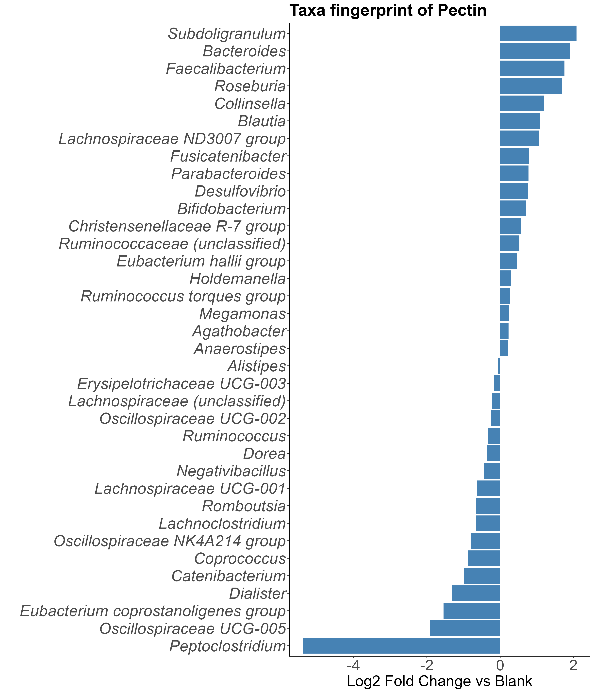

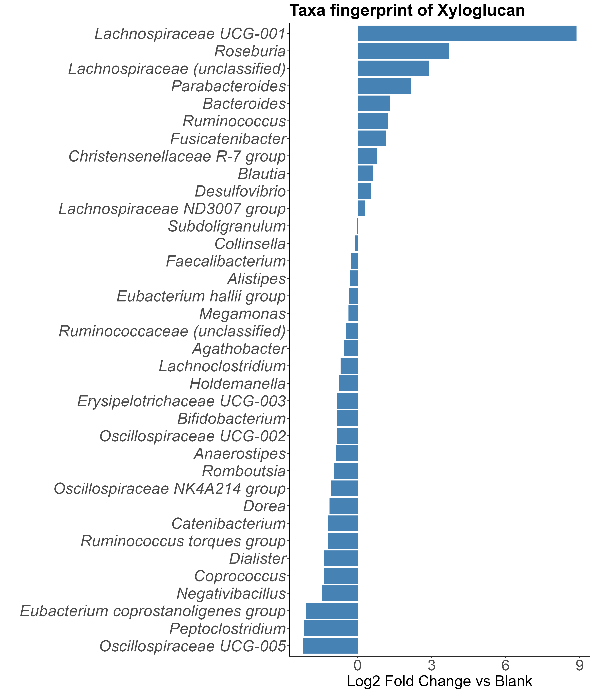

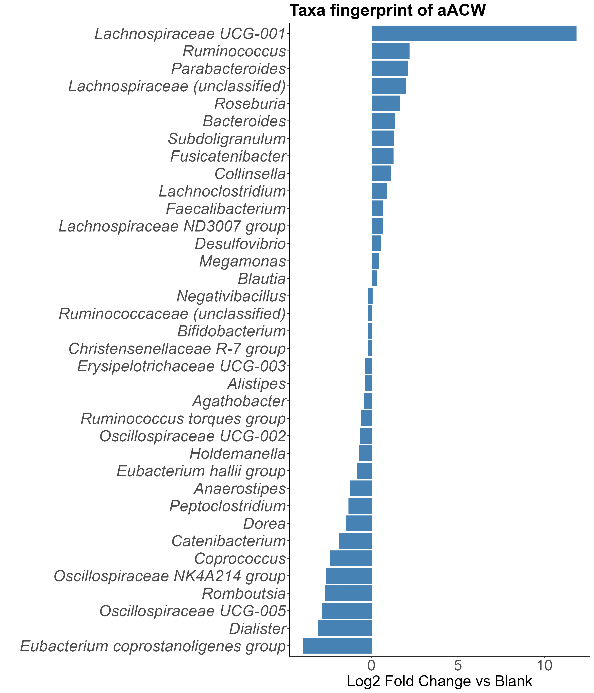

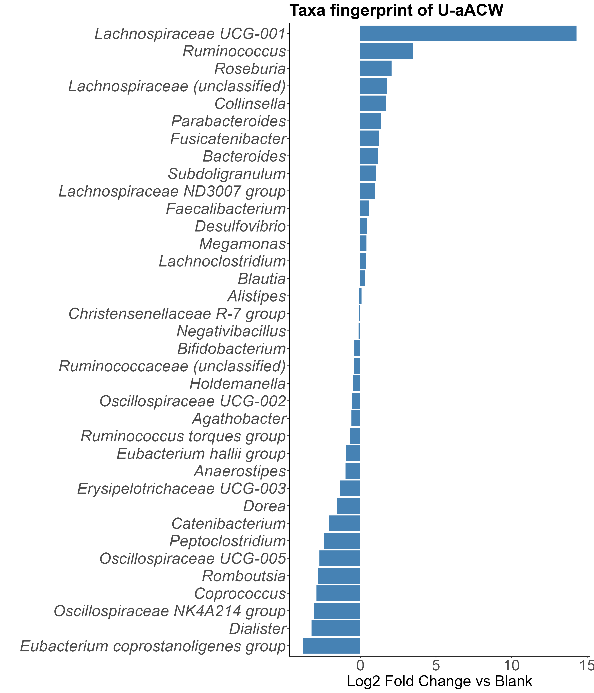

D

G

F

E

C

B

A

Figure S3. Log₂ fold changes in total and individual SCFA and BCFA concentrations following the addition of vacuole-derived phenolic compounds (APP) across substrates (*n = 5*) and time points. Fold changes were calculated from mean concentrations of APP-treated vs. untreated fermentations for the control (fermentation without substrate) and each substrate: total SCFA (A), acetate (B), butyrate (C), propionate (D), total BCFA (E), isovalerate (F) and isobutyrate (G). Error bars represent the bootstrap-estimated SD of the log₂ fold change (1,000 resamplings). Significance between APP-treated and untreated fermentations within each condition and time point was assessed using Welch's two-sample t-test (* p < 0.05, ** p < 0.01, *** p < 0.001; ns, not significant).

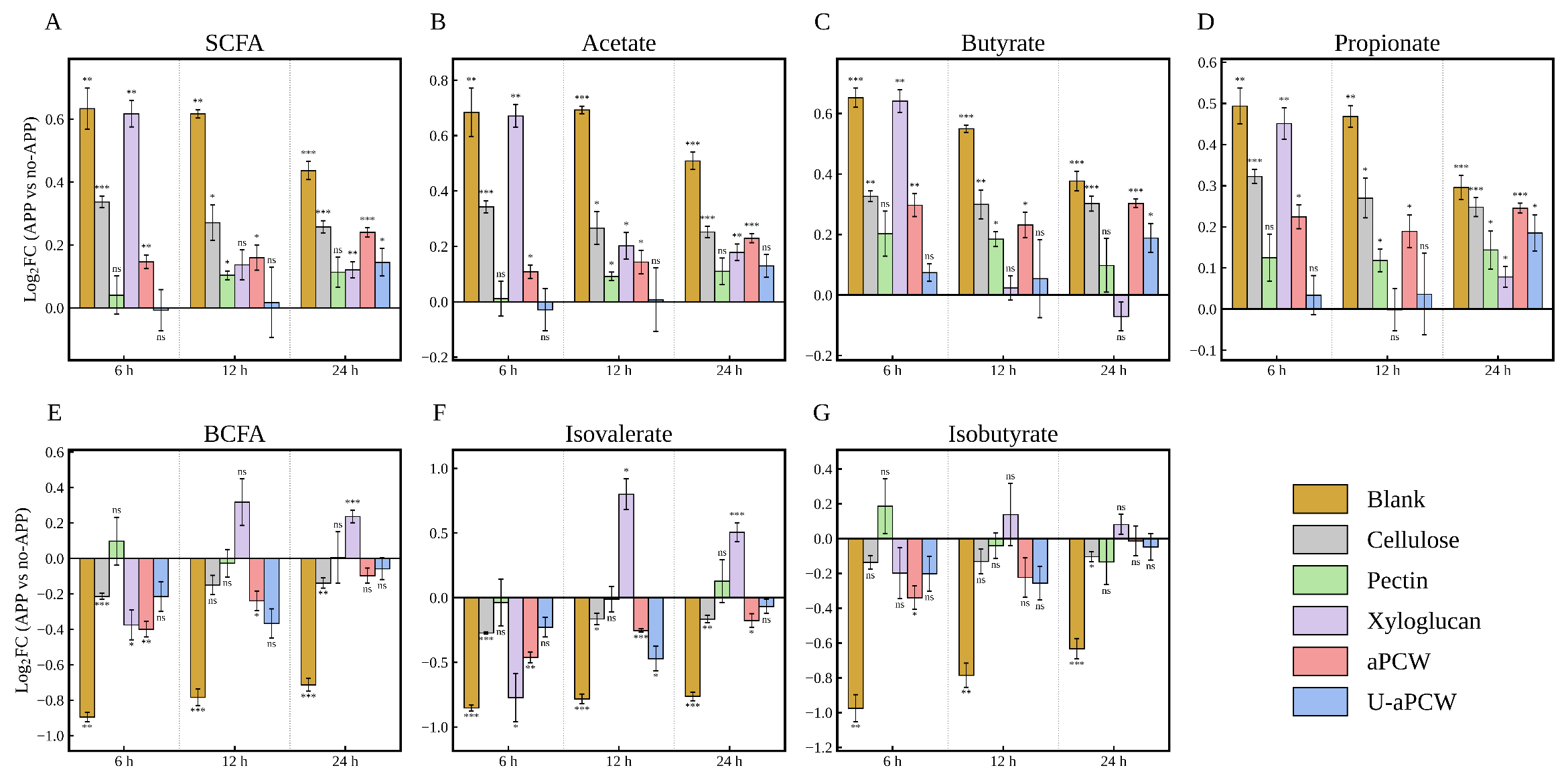

**Table S1.** Concentrations (mM) of short-chain fatty acids (SCFA) and branched-chain fatty acids (BCFA).

| SCFA | Treatment | 6 h | 12 h | 24 h |
| --- | --- | --- | --- | --- |
| **Total SCFA and BCFA pool** | **Control** | 7.49 ± 0.63^a^ | 8.97 ± 0.05^a^ | 10.38 ± 0.22^a^ |
|  | **APP** | 10.81 ± 0.22^a^ | 12.86 ± 0.25^a^ | 13.23 ± 0.59^a^ |
|  | **aPCW** | 40.77 ± 0.80^c^ | 43.44 ± 1.85^b^ | 49.22 ± 0.78^c^ |
|  | **U-aPCW** | 44.58 ± 3.75^c^ | 48.83 ± 7.45^b^ | 47.68 ± 3.17^c^ |
|  | **Cellulose** | 26.74 ± 0.56^b^ | 29.54 ± 2.13^c^ | 33.71 ± 0.97^b^ |
|  | **Xyloglucan** | 28.80 ± 0.73^b^ | 51.04 ± 3.38^bd^ | 65.16 ± 2.47^d^ |
|  | **Pectin** | 67.38 ± 4.38^d^ | 70.35 ± 1.38^e^ | 68.36 ± 4.72^d^ |
|  | **FOS** | 54.44 ± 2.61^e^ | 58.61 ± 0.91^d^ | 67.45 ± 2.33^d^ |
| **Total SCFA** | **Control** | 6.69 ± 0.60^a^ | 8.03 ± 0.02^a^ | 9.28 ± 0.19^a^ |
|  | **APP** | 10.39 ± 0.22^a^ | 12.31 ± 0.25^a^ | 12.56 ± 0.56^a^ |
|  | **aPCW** | 39.96 ± 0.83^c^ | 42.51 ± 1.79^c^ | 47.97 ± 0.79^c^ |
|  | **U-aPCW** | 43.84 ± 3.76^c^ | 47.88 ± 7.46^c^ | 46.54 ± 3.07^c^ |
|  | **Cellulose** | 25.17 ± 0.54^b^ | 27.78 ± 2.01^b^ | 31.65 ± 0.88^b^ |
|  | **Xyloglucan** | 28.56 ± 0.74^b^ | 50.69 ± 3.37^cd^ | 64.37 ± 2.45^d^ |
|  | **Pectin** | 67.14 ± 4.34^d^ | 69.97 ± 1.35^e^ | 67.80 ± 4.66^d^ |
|  | **FOS** | 54.17 ± 2.57^e^ | 58.15 ± 0.92^d^ | 66.56 ± 2.23^d^ |
| **Total BCFA** | **Control** | 0.79 ± 0.03^a^ | 0.94 ± 0.03^a^ | 1.10 ± 0.03^d^ |
|  | **APP** | 0.43 ± 0.00^b^ | 0.55 ± 0.03^b^ | 0.67 ± 0.04^ab^ |
|  | **aPCW** | 0.81 ± 0.03^a^ | 0.93 ± 0.05^a^ | 1.25 ± 0.02^c^ |
|  | **U-aPCW** | 0.74 ± 0.03^a^ | 0.95 ± 0.12^a^ | 1.14 ± 0.10^cd^ |
|  | **Cellulose** | 1.57 ± 0.03^c^ | 1.76 ± 0.12^c^ | 2.06 ± 0.09^e^ |
|  | **Xyloglucan** | 0.24 ± 0.01^d^ | 0.35 ± 0.03^d^ | 0.79 ± 0.04^af^ |
|  | **Pectin** | 0.23 ± 0.04^d^ | 0.38 ± 0.03^bd^ | 0.56 ± 0.08^b^ |
|  | **FOS** | 0.27 ± 0.04^d^ | 0.46 ± 0.03^bd^ | 0.89 ± 0.10^f^ |
| **Acetate** | **Control** | 4.02 ± 0.48^a^ | 4.76 ± 0.04^a^ | 5.49 ± 0.16^a^ |
|  | **APP** | 6.47 ± 0.19^a^ | 7.69 ± 0.14^a^ | 7.81 ± 0.33^a^ |
|  | **aPCW** | 29.71 ± 0.58^c^ | 31.21 ± 1.40^c^ | 34.82 ± 0.69^c^ |
|  | **U-aPCW** | 32.20 ± 3.21^cd^ | 35.03 ± 5.63^c^ | 34.17 ± 2.16^c^ |
|  | **Cellulose** | 17.09 ± 0.44^b^ | 18.97 ± 1.46^b^ | 21.53 ± 0.52^b^ |
|  | **Xyloglucan** | 18.50 ± 0.49^b^ | 32.84 ± 2.15^c^ | 41.73 ± 1.98^d^ |
|  | **Pectin** | 54.10 ± 3.66^e^ | 54.94 ± 1.09^e^ | 52.23 ± 3.61^e^ |
|  | **FOS** | 35.01 ± 1.52^d^ | 37.15 ± 0.76^d^ | 40.48 ± 1.17^d^ |
| **Butyrate** | **Control** | 0.97 ± 0.04^a^ | 1.19 ± 0.01^a^ | 1.36 ± 0.03^a^ |
|  | **APP** | 1.53 ± 0.03^a^ | 1.75 ± 0.02^a^ | 1.77 ± 0.09^a^ |
|  | **aPCW** | 3.68 ± 0.15^b^ | 4.08 ± 0.14^bc^ | 4.86 ± 0.04^c^ |
|  | **U-aPCW** | 4.34 ± 0.14^c^ | 4.87 ± 0.81^c^ | 4.86 ± 0.38^c^ |
|  | **Cellulose** | 3.09 ± 0.01^b^ | 3.33 ± 0.21^b^ | 3.80 ± 0.14^b^ |
|  | **Xyloglucan** | 3.07 ± 0.08^b^ | 7.18 ± 0.42^d^ | 10.85 ± 0.40^e^ |
|  | **Pectin** | 5.00 ± 0.28^d^ | 6.13 ± 0.04^e^ | 6.49 ± 0.47^d^ |
|  | **FOS** | 8.20 ± 0.54^e^ | 8.82 ± 0.04^f^ | 10.95 ± 0.58^e^ |
| **Propionate** | **Control** | 1.70 ± 0.11^a^ | 2.08 ± 0.02^a^ | 2.43 ± 0.04^a^ |
|  | **APP** | 2.39 ± 0.02^a^ | 2.88 ± 0.11^a^ | 2.98 ± 0.14^a^ |
|  | **aPCW** | 6.58 ± 0.14^c^ | 7.22 ± 0.27^c^ | 8.27 ± 0.08^cd^ |
|  | **U-aPCW** | 7.30 ± 0.44^cd^ | 7.98 ± 1.03^cd^ | 7.50 ± 0.53^c^ |
|  | **Cellulose** | 4.99 ± 0.11^b^ | 5.48 ± 0.34^b^ | 6.31 ± 0.22^b^ |
|  | **Xyloglucan** | 6.99 ± 0.20^b^ | 10.67 ± 0.81^e^ | 11.79 ± 0.38^e^ |
|  | **Pectin** | 8.04 ± 0.40^d^ | 8.89 ± 0.29^d^ | 9.08 ± 0.60^d^ |
|  | **FOS** | 10.97 ± 0.53^e^ | 12.18 ± 0.18^f^ | 15.13 ± 0.52^f^ |
| **Isobutyrate** | **Control** | 0.28 ± 0.02^d^ | 0.35 ± 0.03^bd^ | 0.41 ± 0.01^d^ |
|  | **APP** | 0.14 ± 0.01^a^ | 0.20 ± 0.01^a^ | 0.26 ± 0.02^a^ |
|  | **aPCW** | 0.40 ± 0.03^b^ | 0.45 ± 0.05^b^ | 0.59 ± 0.01^b^ |
|  | **U-aPCW** | 0.36 ± 0.02^b^ | 0.44 ± 0.06^b^ | 0.53 ± 0.05^b^ |
|  | **Cellulose** | 0.66 ± 0.03^c^ | 0.76 ± 0.07^c^ | 0.87 ± 0.04^c^ |
|  | **Xyloglucan** | 0.14 ± 0.02^a^ | 0.27 ± 0.03^ad^ | 0.53 ± 0.05^b^ |
|  | **Pectin** | 0.14 ± 0.02^a^ | 0.20 ± 0.02^a^ | 0.28 ± 0.05^a^ |
|  | **FOS** | 0.15 ± 0.03^a^ | 0.25 ± 0.04^ad^ | 0.43 ± 0.06^d^ |
| **Isovalerate** | **Control** | 0.51 ± 0.01^e^ | 0.59 ± 0.01^c^ | 0.69 ± 0.02^c^ |
|  | **APP** | 0.28 ± 0.01^a^ | 0.35 ± 0.02^a^ | 0.41 ± 0.02^a^ |
|  | **aPCW** | 0.41 ± 0.01^b^ | 0.48 ± 0.01^b^ | 0.65 ± 0.02^bc^ |
|  | **U-aPCW** | 0.38 ± 0.02^b^ | 0.51 ± 0.06^bc^ | 0.60 ± 0.04^b^ |
|  | **Cellulose** | 0.91 ± 0.01^c^ | 1.00 ± 0.05^d^ | 1.19 ± 0.05^d^ |
|  | **Xyloglucan** | 0.09 ± 0.02^d^ | 0.08 ± 0.01^e^ | 0.26 ± 0.02^e^ |
|  | **Pectin** | 0.10 ± 0.02^d^ | 0.18 ± 0.01^f^ | 0.29 ± 0.04^e^ |
|  | **FOS** | 0.11 ± 0.01^d^ | 0.59 ± 0.01^f^ | 0.46 ± 0.02^a^ |

Values represent mean ± standard deviation (n=3). Within each SCFA type and time point, values with different letters are significantly different (p < 0.05) for each SCFA and time point combination according to Tukey's HSD test.

**Table S2.** Pearson correlation coefficients (r) and corresponding p-values for associations between gut microbial taxa and individual microbial metabolites (SCFAs and BCFAs).

|  | **Acetate** | | **Butyrate** | | **Propionate** | | **Isobutyrate** | | **Isovalerate** | |
| --- | --- | --- | --- | --- | --- | --- | --- | --- | --- | --- |
|  | **r** | ***p*** | **r** | ***p*** | **r** | ***p*** | **r** | ***p*** | **r** | ***p*** |
| *Agathobacter* | -0.060 | 0.778 | 0.026 | 0.910 | 0.092 | 0.679 | -0.245 | 0.215 | 0.060 | 0.778 |
| *Alistipes* | -0.304 | 0.120 | -0.461 | 0.011 | -0.443 | 0.015 | -0.167 | 0.403 | 0.128 | 0.538 |
| *Anaerostipes* | 0.039 | 0.863 | 0.209 | 0.299 | 0.256 | 0.199 | -0.327 | 0.088 | -0.127 | 0.539 |
| *Bacteroides* | 0.793 | <0.001 | 0.796 | <0.001 | 0.783 | <0.001 | -0.260 | 0.193 | -0.552 | 0.001 |
| *Bifidobacterium* | 0.169 | 0.399 | 0.070 | 0.759 | 0.151 | 0.456 | -0.379 | 0.045 | -0.082 | 0.717 |
| *Blautia* | 0.426 | 0.020 | 0.582 | <0.001 | 0.568 | <0.001 | -0.521 | 0.003 | -0.607 | <0.001 |
| *Catenibacterium* | -0.773 | <0.001 | -0.697 | <0.001 | -0.692 | <0.001 | -0.227 | 0.251 | 0.241 | 0.220 |
| *Christensenellaceae R-7 group* | 0.359 | 0.059 | 0.360 | 0.059 | 0.245 | 0.215 | -0.254 | 0.202 | -0.590 | <0.001 |
| *Collinsella* | 0.403 | 0.030 | 0.170 | 0.399 | 0.283 | 0.152 | 0.121 | 0.559 | 0.273 | 0.168 |
| *Coprococcus* | -0.841 | <0.001 | -0.843 | <0.001 | -0.863 | <0.001 | -0.311 | 0.110 | 0.150 | 0.459 |
| *Desulfovibrio* | 0.729 | <0.001 | 0.661 | <0.001 | 0.649 | <0.001 | -0.164 | 0.411 | -0.460 | 0.011 |
| *Dialister* | -0.749 | <0.001 | -0.638 | <0.001 | -0.667 | <0.001 | -0.170 | 0.399 | 0.183 | 0.367 |
| *Dorea* | -0.554 | 0.001 | -0.681 | <0.001 | -0.629 | <0.001 | 0.108 | 0.615 | 0.577 | <0.001 |
| *Erysipelotrichaceae UCG-003* | -0.408 | 0.028 | -0.371 | 0.050 | -0.297 | 0.128 | -0.175 | 0.390 | 0.250 | 0.211 |
| *Eubacterium coprostanoligenes group* | -0.855 | <0.001 | -0.831 | <0.001 | -0.848 | <0.001 | -0.243 | 0.217 | 0.246 | 0.215 |
| *Eubacterium hallii group* | 0.034 | 0.874 | 0.182 | 0.368 | 0.190 | 0.348 | -0.575 | <0.001 | -0.383 | 0.043 |
| *Faecalibacterium* | 0.454 | 0.013 | 0.282 | 0.152 | 0.333 | 0.084 | -0.456 | 0.012 | -0.332 | 0.084 |
| *Fusicatenibacter* | 0.795 | <0.001 | 0.817 | <0.001 | 0.769 | <0.001 | -0.069 | 0.763 | -0.539 | 0.002 |
| *Holdemanella* | -0.171 | 0.399 | -0.339 | 0.080 | -0.274 | 0.167 | -0.075 | 0.742 | 0.299 | 0.126 |
| *Lachnoclostridium* | -0.371 | 0.050 | -0.447 | 0.014 | -0.351 | 0.066 | 0.258 | 0.196 | 0.522 | 0.003 |
| *Lachnospiraceae (unclassified)* | 0.203 | 0.314 | 0.232 | 0.242 | 0.164 | 0.411 | 0.442 | 0.016 | 0.016 | 0.939 |
| *Lachnospiraceae ND3007 group* | 0.511 | 0.004 | 0.570 | <0.001 | 0.607 | <0.001 | -0.379 | 0.045 | -0.407 | 0.028 |
| *Lachnospiraceae UCG-001* | 0.248 | 0.214 | 0.245 | 0.215 | 0.187 | 0.356 | 0.224 | 0.258 | -0.183 | 0.367 |
| *Megamonas* | -0.106 | 0.620 | -0.024 | 0.914 | 0.041 | 0.856 | -0.364 | 0.056 | -0.066 | 0.771 |
| *Negativibacillus* | -0.084 | 0.712 | -0.307 | 0.115 | -0.191 | 0.348 | 0.592 | <0.001 | 0.834 | <0.001 |
| *Oscillospiraceae NK4A214 group* | -0.680 | <0.001 | -0.519 | 0.003 | -0.559 | <0.001 | -0.436 | 0.017 | -0.062 | 0.778 |
| *Oscillospiraceae UCG-002* | -0.486 | 0.007 | -0.669 | <0.001 | -0.615 | <0.001 | 0.054 | 0.802 | 0.482 | 0.007 |
| *Oscillospiraceae UCG-005* | -0.832 | <0.001 | -0.849 | <0.001 | -0.838 | <0.001 | 0.009 | 0.968 | 0.443 | 0.015 |
| *Parabacteroides* | 0.707 | <0.001 | 0.880 | <0.001 | 0.824 | <0.001 | 0.241 | 0.220 | -0.334 | 0.083 |
| *Peptoclostridium* | -0.411 | 0.027 | -0.335 | 0.082 | -0.295 | 0.131 | 0.617 | <0.001 | 0.677 | <0.001 |
| *Romboutsia* | -0.669 | <0.001 | -0.550 | 0.001 | -0.606 | <0.001 | -0.494 | 0.005 | -0.148 | 0.462 |
| *Roseburia* | 0.653 | <0.001 | 0.790 | <0.001 | 0.687 | <0.001 | -0.063 | 0.778 | -0.620 | <0.001 |
| *Ruminococcaceae (unclassified)* | 0.038 | 0.863 | 0.080 | 0.724 | 0.130 | 0.534 | -0.553 | 0.001 | -0.272 | 0.168 |
| *Ruminococcus* | 0.330 | 0.086 | 0.227 | 0.251 | 0.205 | 0.311 | 0.336 | 0.082 | -0.004 | 0.985 |
| *Ruminococcus torques group* | -0.372 | 0.050 | -0.499 | 0.005 | -0.440 | 0.016 | -0.426 | 0.020 | 0.061 | 0.778 |
| *Subdoligranulum* | 0.329 | 0.087 | -0.023 | 0.915 | 0.002 | 0.990 | -0.286 | 0.146 | -0.202 | 0.315 |

**Table S3.** Shrunken log₂ fold changes of microbial taxa in APP-treated versus untreated fermentations across different fiber substrates (DESeq2 analysis).

| **Taxa** | **Cellulose** | **aPCW** | **U-aPCW** | **Pectin** | **Xyloglucan** |
| --- | --- | --- | --- | --- | --- |
| *Agathobacter* | -0.094 | 0.018 | 0.068 | 0.000 | 0.027 |
| *Alistipes* | 0.062 | -0.006 | -0.042 | 0.001 | 0.032 |
| *Anaerostipes* | 0.085 | -0.008 | 0.079 | 0.002 | 0.102 |
| *Bacteroides* | 0.003 | 0.000 | -0.132 | -0.001 | **-0.346** |
| *Bifidobacterium* | -0.015 | -0.013 | 0.012 | 0.001 | 0.123 |
| *Blautia* | **0.190*** | 0.014 | 0.000 | 0.000 | 0.072 |
| *Catenibacterium* | 0.062 | 0.024 | 0.011 | 0.003 | -0.034 |
| *Christensenellaceae R-7 group* | 0.036 | -0.010 | -0.018 | -0.002 | -0.085 |
| *Collinsella* | -0.038 | -0.003 | -0.005 | 0.001 | 0.072 |
| *Coprococcus* | -0.009 | -0.015 | 0.023 | -0.066 | -0.191 |
| *Desulfovibrio* | 0.044 | -0.016 | -0.032 | -0.002 | -0.015 |
| *Dialister* | -0.014 | -0.016 | 0.010 | 0.001 | -0.015 |
| *Dorea* | **-0.315** | -0.007 | -0.006 | -0.001 | -0.038 |
| *Erysipelotrichaceae UCG-003* | 0.072 | -0.009 | 0.011 | -0.002 | -0.045 |
| *Eubacterium coprostanoligenes group* | -0.147 | -0.003 | 0.007 | 0.002 | -0.040 |
| *Eubacterium hallii group* | -0.020 | 0.033 | 0.016 | -0.001 | 0.059 |
| *Faecalibacterium* | -0.136 | **0.169** | 0.005 | 0.001 | 0.043 |
| *Fusicatenibacter* | 0.020 | -0.001 | -0.040 | -0.001 | **0.389** |
| *Holdemanella* | -0.157 | 0.018 | 0.052 | -0.002 | 0.005 |
| *Lachnoclostridium* | -0.081 | -0.013 | 0.001 | **1.013** | 0.066 |
| *Lachnospiraceae (unclassified)* | 0.028 | -0.005 | 0.008 | 0.027 | **-0.542** |
| *Lachnospiraceae ND3007 group* | 0.009 | 0.000 | 0.001 | -0.013 | 0.032 |
| *Lachnospiraceae UCG-001* | 0.000 | -0.009 | -0.090 | 0.000 | **-0.547** |
| *Megamonas* | 0.208 | -0.001 | 0.164 | 0.000 | -0.226 |
| *Negativibacillus* | -0.187 | 0.029 | 0.001 | -0.006 | 0.020 |
| *Oscillospiraceae NK4A214 group* | -0.072 | 0.009 | -0.012 | -0.001 | 0.032 |
| *Oscillospiraceae UCG-002* | 0.009 | 0.009 | -0.019 | -0.002 | 0.109 |
| *Oscillospiraceae UCG-005* | 0.022 | -0.001 | 0.040 | 0.021 | -0.010 |
| *Parabacteroides* | -0.017 | -0.002 | -0.093 | 0.008 | -0.146 |
| *Peptoclostridium* | -0.226 | -0.013 | -0.013 | -0.001 | **-0.397** |
| *Romboutsia* | -0.181 | 0.000 | -0.031 | 0.001 | -0.112 |
| *Roseburia* | 0.156 | 0.006 | -0.006 | -0.008 | **-0.510** |
| *Ruminococcaceae (unclassified)* | **0.361** | 0.054 | 0.021 | 0.000 | 0.064 |
| *Ruminococcus* | -0.003 | -0.008 | **-0.250** | -0.001 | 0.018 |
| *Ruminococcus torques group* | 0.144 | 0.007 | **0.228** | 0.000 | **0.443** |
| *Subdoligranulum* | -0.079 | 0.004 | -0.007 | -0.015 | 0.002 |

Values in bold are those 14 changes with a significant log2FC (p<0.05).
